# Reconciling the potentially irreconcilable? Genotypic and phenotypic amoxicillin-clavulanate resistance in *Escherichia coli*

**DOI:** 10.1101/511402

**Authors:** Timothy J. Davies, Nicole Stoesser, Anna E Sheppard, Manal Abuoun, Philip Fowler, Jeremy Swann, T. Phuong Quan, David Griffiths, Alison Vaughan, Marcus Morgan, Hang TT Phan, Katie J Jeffery, Monique Andersson, Matt J Ellington, Oskar Ekelund, Neil Woodford, Amy J. Mathers, Robert A. Bonomo, Derrick W. Crook, Tim E.A. Peto, Muna F Anjum, A. Sarah Walker

## Abstract

Resistance to amoxicillin-clavulanate, a widely used beta-lactam/beta-lactamase inhibitor combination antibiotic, is rising globally, yet susceptibility testing remains challenging. To test whether whole-genome sequencing (WGS) could provide a more reliable assessment of susceptibility than traditional methods, we predicted resistance from WGS for 976 *E. coli* bloodstream infection isolates from Oxfordshire, UK, comparing against phenotypes from the BD Phoenix (calibrated against EUCAST guidelines). 339/976 (35%) isolates were amoxicillin-clavulanate resistant. Predictions based solely on beta-lactamase presence/absence performed poorly (sensitivity 23% (78/339)) but improved when genetic features associated with penicillinase hyper-production (e.g. promoter mutations, copy number estimates) were considered (sensitivity 82% (277/339); p<0.0001). Most discrepancies occurred in isolates with peri-breakpoint MICs. We investigated two potential causes; the phenotypic reference and the binary resistant/susceptible classification. We performed reference standard, replicated phenotyping in a random stratified subsample of 261/976 (27%) isolates using agar dilution, following both EUCAST and CLSI guidelines, which use different clavulanate concentrations. As well as disagreeing with each other, neither agar dilution phenotype aligned perfectly with genetic features. A random-effects model investigating associations between genetic features and MICs showed that some genetic features had small, variable and additive effects, resulting in variable resistance classification. Using model fixed-effects to predict MICs for the non-agar dilution isolates, predicted MICs were in essential agreement (±1 doubling dilution) with observed (BD Phoenix) MICs for 691/715 (97%) isolates. This suggests amoxicillin-clavulanate resistance in *E. coli* is quantitative, rather than qualitative, explaining the poorly reproducible binary (resistant/susceptible) phenotypes and suboptimal concordance between different phenotypic methods and with WGS-based predictions.

## Introduction

Rising amoxicillin-clavulanate resistance in *E. coli* is a major healthcare challenge, with increasing incidence of resistant bloodstream infections (BSI)(1) threatening its utility as the most commonly used antibiotic in Europe.(2) Consequently, many hospitals are considering broadening their first-line empiric antibiotics for common infections. However, significant uncertainty is created by observed differences between the two main assays for amoxicillin-clavulanate susceptibility in the classification of clinical samples.(3) These differences are so large that increasing amoxicillin-clavulanate resistance was suggested to be primarily due to laboratories switching from US Clinical Laboratory Standards Institute (CLSI) to European Committee on Antimicrobial Susceptibility Testing (EUCAST) guidelines.(4) Recent work,(5) however, suggests that changes in laboratory protocols are unlikely to account for the majority of the increase in resistance. Only one study has investigated whether there are underlying genetic causes for the ongoing rise in amoxicillin-clavulanate resistance,(6) but found no evidence of clonal expansion of any specific amoxicillin-clavulanate-resistant strains. However, the genetic epidemiology of amoxicillin-clavulanate resistance mechanisms was not investigated.

In addition to its widespread clinical use, amoxicillin-clavulanate is a model for beta-lactam/beta-lactamase inhibitor (BL/BLI) combinations, which are the focus of renewed attention(7) due to the development of novel BL/BLIs with activity against highly drug-resistant organisms.(8) EUCAST has recently published guidelines on setting breakpoints for BL/BLIs,(9) but the inconsistencies seen in testing and clinically interpreting amoxicillin-clavulanate resistance likely extend to novel BL/BLIs.(10)

One solution is to instead identify the genetic determinants characterizing resistance (resistance genotype) using whole-genome sequencing (WGS).(11) This approach may be particularly helpful for BL/BLI, as recent studies have suggested that traditional phenotyping is less accurate in isolates producing extended spectrum beta-lactamases.(12) Rather than resistance being associated with the simple presence/absence of specific genes, previous studies have found much amoxicillin-clavulanate resistance is likely attributable to mechanisms which increase the effective concentration of beta-lactamases (e.g. additive effects of multiple beta-lactamases,(13) increasing gene expression(14) or modifying cell permeability(15)). Given the added complexity of both phenotype and genotype, studies using WGS to predict phenotypic resistance have either not included amoxicillin-clavulanate,(16, 17) compared against only one set of breakpoints,(18) or only tested small sets of pre-selected samples.(19) Similar studies investigating other BL/BLIs, such as piperacillin-tazobactam, reported poor accuracy when predicting resistance from genotype.(20)

We therefore investigated concordance between WGS-derived genotypes and amoxicillin-clavulanate susceptibility phenotypes in a large, unselected set of Oxfordshire *E. coli* BSI isolates from 2013-2015. We assessed whether extending the usual presence/absence genetic approach to include features that might increase beta-lactamase expression (copy number, promoter type) would improve concordance, and quantified the impact of particular genetic variants and testing guidelines (EUCAST, CLSI) on minimum inhibitory concentrations (MICs).

## Results

### Routine laboratory phenotypes and amoxicillin-clavulanate resistance genotypes

Of the 1039 *E. coli* BSI occurring between January 2013-August 2015 in Oxfordshire, UK, 1000 had at least one isolate stored by Oxford University Hospitals (OUH) NHS Foundation Trust microbiology laboratory. In most (992/1000 (99%)) infections, only a single *E. coli* was isolated; however, two different *E. coli* were grown from culture in 8 cases, giving a total of 1008 distinct *E. coli* isolates. Each of these isolates had linked antimicrobial susceptibility test (AST) data from the Oxford University Hospitals (OUH) NHS Foundation Trust microbiology laboratory using the BD Phoenix (Beckton, Dickinson and Company). All obtained isolates were sequenced, with 976/1008 (97%) having WGS data meeting pre-determined quality controls designed to identify mixtures and poor-quality sequences. Overall, these 976 isolates represented 968/1039 [93%] *E. coli* BSI (Supplementary Figure S1). 339/976 (36%) had amoxicillin-clavulanate MIC > 8/2 mg/L by EUCAST breakpoints (Supplementary Table S1).

The collection was highly diverse, representing 152 different sequence types (STs). The most common was ST73 (161,17%) (Supplementary Figure S2), followed by ST131 (124,13%), which had the highest percentage of phenotypically-resistant isolates (N=74,60%) and was the only ST associated with amoxicillin-clavulanate resistance (chi-squared p<0.0001 compared with p>0.16 for all other STs).

The most common beta-lactam resistance mechanisms identified (using ARIBA(21) (default parameters) and tBLASTn/BLASTn (see Methods)) were acquired beta-lactamase genes, which were identified in 515/976 (53%) isolates. Most of these (448/515 (87%)) harbored only a single transmissible beta-lactamase gene. Among the 67 isolates with more than one beta-lactamase gene, the most common combination was *bla*_CTX-M-15_ and *bla*_OXA-1_ (N=27, Supplementary Table S2B). Overall *bla*_TEM_ was by far the most common mechanism identified (occurring in 427/976 (44%) isolates), followed by *bla*_CTX-M_ (N=73 (7%)), *bla*_OXA_ (N=62 (6%)) and *bla*_SHV_ (N=23 (2%)) (Supplementary Figure S2, Supplementary Table S2). For the 594 transmissible beta-lactamases identified, median DNA copy number from mapping coverage was 2.23 (IQR 1.73,3.31); 227 (38%) had >2.5-fold coverage (the threshold to predict resistance derived from receiver operating characteristic (ROC) analysis of isolates with only one beta-lactamase identified, Supplementary Methods, Supplementary Figure S3). Variant *bla*_TEM_ and *ampC* promoters considered to be associated with increased expression were identified in 49 (5%) and 20 (2%) isolates respectively (Supplementary Table 3A, 3C). 31 (3%) isolates potentially had one non-functional porin, of which 22 also contained a beta-lactamase gene; however, no isolate had “functionally lost” both *ompC* and *ompF* (Supplementary Table S4).

### WGS-derived resistance prediction compared with routine phenotyping

We compared two genetic resistance prediction algorithms for amoxicillin and amoxicillin-clavulanate (see Methods). The first, denoted the “basic” prediction algorithm, was analogous to common WGS-based resistance methods and only predicting resistance for isolates containing inhibitor resistant beta-lactamase genes (e.g. *bla*_OXA-1_, *bla*_TEM-30_). The second denoted the “extended” prediction algorithm, additionally evaluated *bla*_TEM_ and *ampC* promoter mutations, estimates of beta-lactamase gene DNA copy number and porin loss-of-function mutations. Including the additional features (i.e. the “extended” approach) had little impact on our ability to identify ampicillin resistance but significantly improved amoxicillin-clavulanate resistance prediction (Table 1, amoxicillin sensitivity 98% basic vs 96% extended; amoxicillin-clavulanate sensitivity 23% basic vs 82% extended, McNemar’s p<0.0001). However, the increased sensitivity also came at the cost of modestly reduced specificity (Table 1). Overall categorical agreement of WGS-derived with observed phenotype increased from 712 (73%) to 868 (89%) when these extended genetic features were included.

**Table 1:** Performance of WGS-based prediction using both basic and extended algorithms

| **WGS-predicted phenotype**<br>**(No. isolates)** | **AST Phenotype**<br>**(No. isolates)** |  | **Sensitivity**<br>**(%)** | **Specificity**<br>**(%)** |
| --- | --- | --- | --- | --- |
| **Ampicillin: basic** | S | R | 96 | 99 |
| S | 441 | 20 |  |  |
| R | 4 | 511 |  |  |
| **Ampicillin: extended prediction** | S | R | 98 | 99 |
| S | 439 | 8 |  |  |
| R | 6 | 523 |  |  |
| **Amoxicillin-clavulanate: basic** | S | R | 23 | 100 |
| S | 634 | 261 |  |  |
| R | 3 | 78 |  |  |
| **Amoxicillin-clavulanate: extended** | S | R | 82 | 93 |
| S | 591 | 62 |  |  |
| R | 46 | 277 |  |  |

Investigating the cause of lower than optimal agreement, even using the extended algorithm, showed that most false positive predictions were made on the basis of increased beta-lactamase gene DNA copy-number (**Error! Reference source not found.**). Although there was a clear association between increasing copy-number and MIC (p<0.0001), resistance prediction based on increased DNA copy-number (>2.5) was less accurate than other extended algorithm components (positive predictive value (PPV) = 0.77 compared to >0.97 for all other algorithm components), with both resistant isolates with lower copy-number beta-lactamases and susceptible isolates with higher copy beta-lactamases (Figure 1).

**Figure 1:**
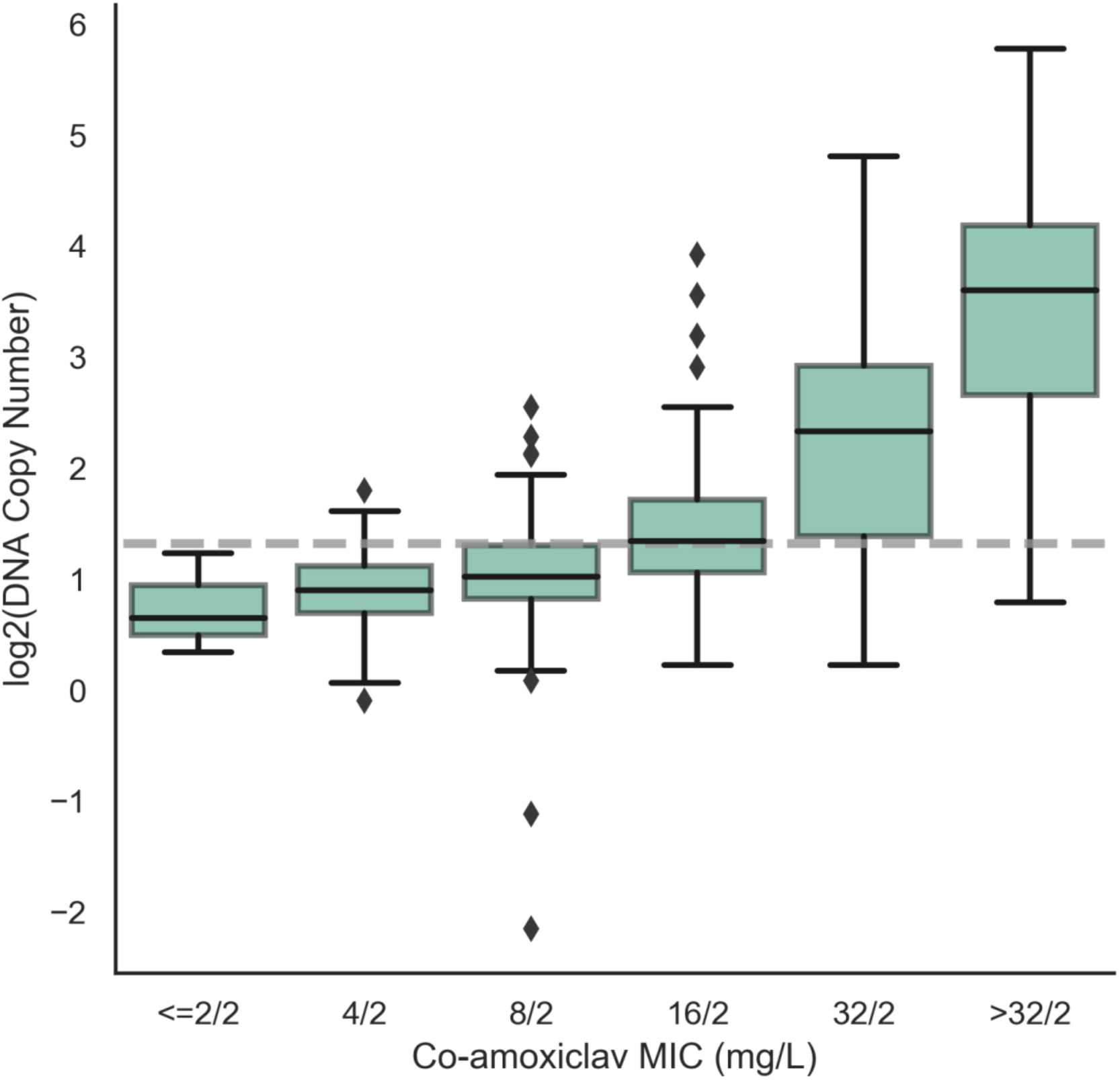
Association between transmissible beta-lactamase gene DNA copy number and amoxicillin-clavulanate MIC in isolates with no alternative resistance features. Note: Evidence for association between MIC and log2(DNA copy number) p<0.0001, estimated using quantile regression. Grey line indicates 2.5 threshold used to define resistance in the extended algorithm based on ROC analysis (Supplementary Figure S3A). Of these 328 isolates, 294 had *bla*_TEM_ genes (290 with *bla*_TEM-1_, 4 with other non-inhibitor resistant *bla*_TEM_ genes), 19 had non-inhibitor resistant *bla*_SHV_ genes and 15 had *bla*_CTX-M_ genes.

The distribution of MICs in isolates with concordant vs discordant predictions suggested an alternative explanation (Figure 2), with the extended algorithm performing better at predicting susceptibility/resistance in non-peri-breakpoint isolates. Overall the algorithm correctly classified 463/469 (99%) isolates with MIC <=4/2 mg/L as susceptible and 230/250 (92%) isolates with MIC >=32/2 mg/L as resistant. Notably, of 79 discordant isolates containing only non-inhibitor-resistant beta-lactamases, 64 (81%) had peri-breakpoint (8/2-16/2mg/L) MICs. Given these findings, we therefore investigated two other hypotheses that could explain the low agreement in peri-breakpoint isolates: (*i*) variable accuracy of the different phenotypic methods, and (*ii*) the binary resistant/susceptible classification being too simplistic.

**Figure 2:**
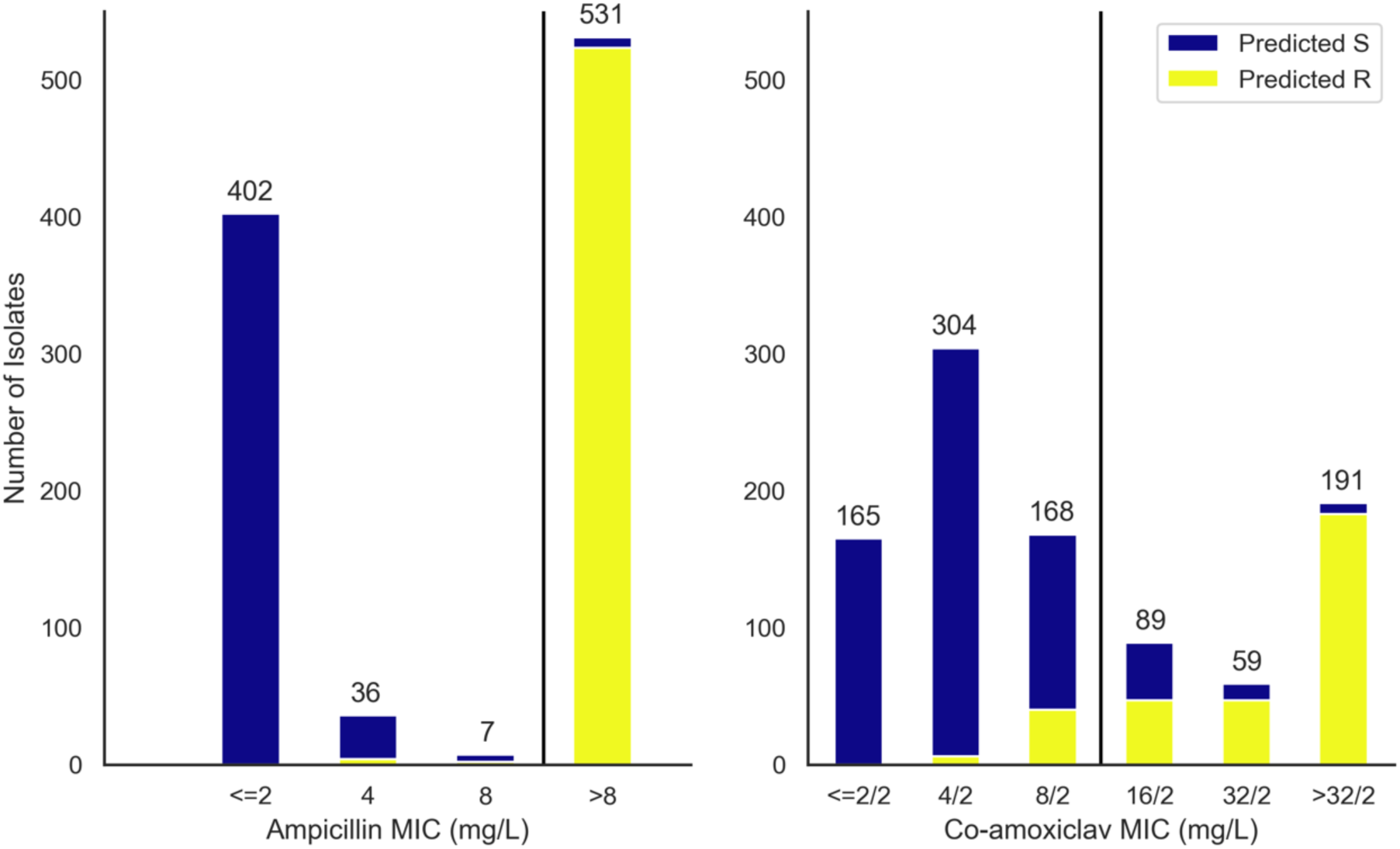
Proportion WGS predicted resistant (extended algorithm) by routine laboratory MIC

### Variability in reference standard agar dilution phenotypes (EUCAST and CLSI based)

291/976 (30%) isolates were selected for repeated agar dilution phenotyping using stratified random subsampling to enrich for resistant isolates both with and without beta-lactamase genes (see Methods, Supplementary Methods, Supplementary Figure S1). Of these 291 isolates, 261 (90%) passed the additional quality control steps designed to remove potential undetected mixtures, and were included in the agar-dilution subsample (details in Supplementary Methods; in brief, all colonies had to be of one morphology on blood-agar purity plates and MICs for each of amoxicillin and both amoxicillin-clavulanate fixed (EUCAST)/ratio (CLSI) tests had to be in essential agreement on 2 or more repeat tests). The stratified random sampling enriching for resistant phenotypes meant that 160/261 (61%) subsample isolates were amoxicillin-clavulanate-resistant by routine AST (Supplementary Table S1). All STs with >10 isolates in the main sample were represented, with 52 (20%), 43 (16%) and 29 (11%) isolates being ST131, ST73 and ST69, respectively, as were all resistance gene families in the main sample (Supplementary Figure S2).

As expected, phenotypes from different reference-standard AST methods were often discordant (Figure 3). EUCAST-based agar dilution (using the fixed 2mg/L clavulanate concentration) only agreed with CLSI-based agar dilution (using the 2:1 ratio of amoxicillin:clavulanate) for 143/261 (55%) isolates (27 agreed resistant, 116 agreed susceptible). For the remaining 118 isolates, EUCAST-based agar dilution results were more conservative than CLSI-based agar dilution. Major discrepancies occurred for 39 isolates, being classed resistant by EUCAST-based agar dilution and susceptible by CLSI-based agar dilution. The remaining 79 isolates were EUCAST-resistant CLSI-intermediate. Excluding isolates classified as intermediate by CLSI, categorical agreement between the two reference-standard methods was 79%. Considering CLSI-intermediate as resistant had little impact on the overall categorical agreement (85%). Each of these test methods also often classified isolates differently to the BD Phoenix (Figure 3), but, as expected, given the BD Phoenix used in the OUH routine laboratory is calibrated against EUCAST guidelines, EUCAST-based agar dilution was in agreement more often. Of note, one isolate which only contained a partial *bla*_TEM_ gene was repeatedly resistant on both EUCAST and CLSI-based agar dilution testing but was identified susceptible by the BD Phoenix.

**Figure 3:**
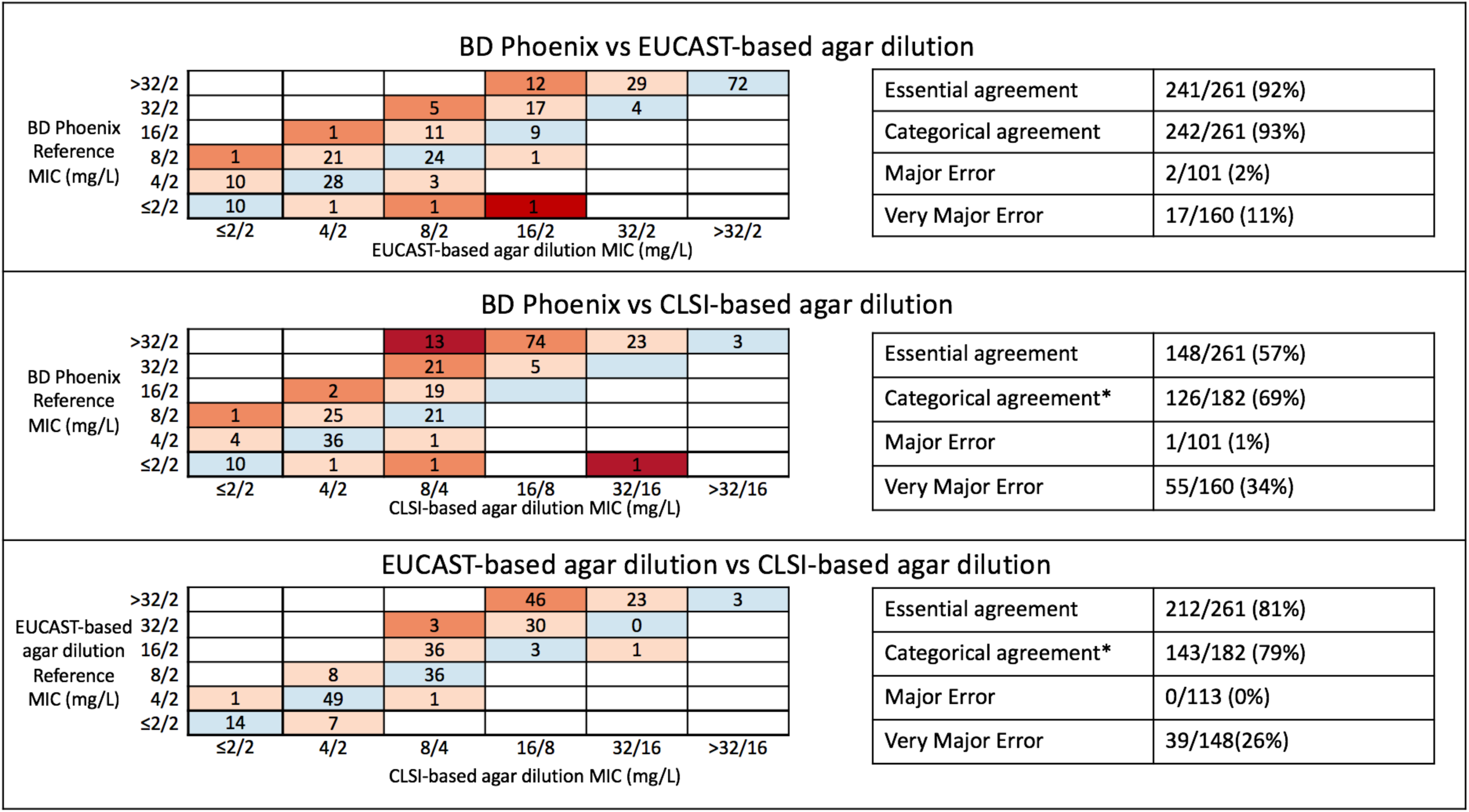
Comparison of the three different phenotyping methods on the agar-dilution subsample isolates (N=261) Note: Comparison of MICs obtained using the three different phenotyping methods; EUCAST-based agar-dilution, CLSI-based agar dilution and BD Phoenix (performed in the OUH microbiology laboratory and using panels calibrated against EUCAST guidelines). *: Isolates are in categorical agreement if they are reported as either resistant by both methods (i.e. BD Phoenix/EUCAST-based agar dilution MIC >8/2 mg/L and CLSI-based agar dilution > 16/8 mg/L) or susceptible by both methods (i.e. BD Phoenix/EUCAST-based agar dilution MIC <=8/2 mg/L and CLSI-based agar dilution MIC <= 8/4 mg/L). Intermediate isolates were excluded from these comparisons (but are shown above) as BD phoenix/EUCAST-based agar dilution have no intermediate category. Blue: full agreement of MICs, light orange: essential agreement, dark orange: within two doubling dilutions (theoretically feasible believing both tests having an error of +/- 1 dilution) and red: disagreement.

MIC results from both methods were variable on retesting (as part of triplicate repeats): more so for EUCAST-based agar dilution MICs (Supplementary Figure S4), which were not constant across repeats for 158/261 (61%) isolates versus only 73 (28%) for CLSI-based agar dilution MICs. While differences across repeats were in essential agreement with one another (i.e. less than (±1 doubling dilution) for all but 12 isolates for EUCAST-based agar dilution and all but 1 isolate for CLSI-based agar dilution, they did cause changes in resistance classification. For EUCAST-based agar dilution, 40/261 (15%) isolates were identified as both resistant and susceptible across repeats, suggesting that even within-method categorical agreement is far poorer than the standards required for regulatory approval. Likewise for CLSI-based agar dilution, MIC differences across repeats resulted in variation in resistance classification for 31 (12%) isolates; however because of the CLSI-intermediate category, 28/31 (90%) of these would be classed as minor discrepancies.

### WGS-derived resistance prediction compared with reference-standard agar dilution phenotypes

Overall, using the extended algorithm above, WGS classified as resistant 23/27 (85%) isolates agreed resistant by EUCAST and CLSI-based tests, 107/118 (91%) indeterminate isolates (76/79 EUCAST-resistant/CLSI-intermediate, 31/39 EUCAST-resistant/CLSI-susceptible) and 17/116 (15%) agreed susceptible isolates (Figure 4). Again predictions based on the presence of high copy number (>2.5x) non-inhibitor beta-lactamases alone were the least congruent with the reference-standard phenotypes. Specifically, 16/62 (26%) isolates with increased copy number beta-lactamase genes were agreed susceptible (accounting for 16 of the 17 resistance predictions in agreed susceptible isolates). Further, whilst 46/62 (74%) isolates with this mechanism were resistant on EUCAST-based agar dilution, similar to the PPV with BD Phoenix on the whole dataset, for CLSI-based agar dilution only 2/62 (3%) were CLSI resistant, 24/62 (39%) were CLSI-intermediate and 36/62 (58%) were CLSI susceptible, suggesting this threshold performs more poorly for predicting CLSI-based agar dilution phenotypes. However, as when selecting the initial threshold against BD Phoenix results, there was no threshold which perfectly predicted the CLSI-based phenotype (Supplementary Figure S2B). Only 8 (30%) of the 27 agreed resistant by EUCAST and CLSI-based tests contained inhibitor-resistant beta-lactamases. Conversely, 24/79 (30%) CLSI-intermediate and 10/39 (26%) CLSI-susceptible isolates contained *bla*_OXA-1_, showing that identification of inhibitor-resistant beta-lactamases was neither necessary nor sufficient to predict resistance for the CLSI-based tests.

**Figure 4.**
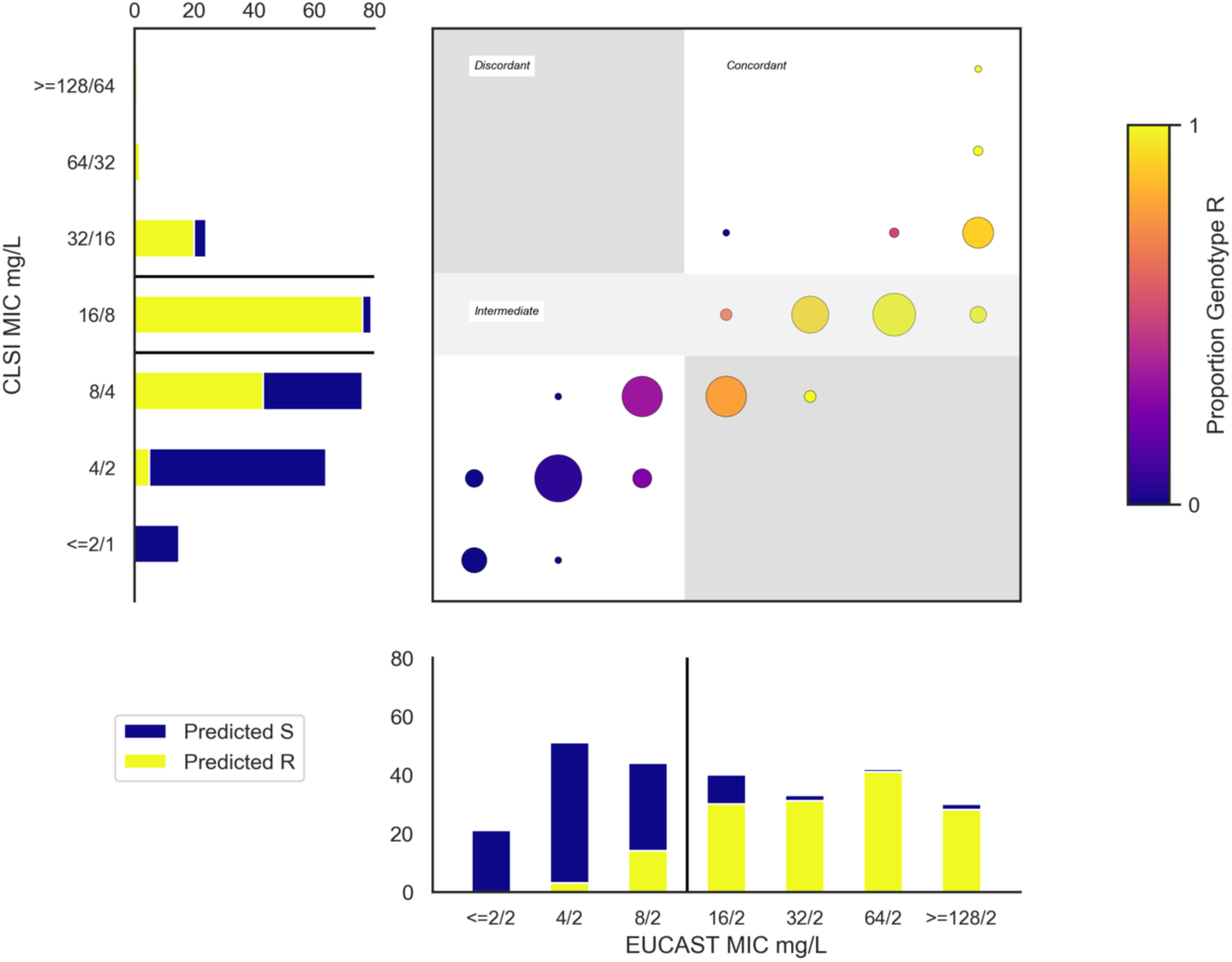
Proportion WGS predicted resistant (extended algorithm) by MICs from EUCAST and CLSI-based methods. Note: Main panel, each (x,y) coordinate represents (EUCAST-based MIC,CLSI-based MIC) combination. At each coordinate, circle size represents the number of isolates with this combination of fixed and ratio MICs, and color denotes proportion identified as resistant by WGS as indicated by the color bar to the right of the figure. The two sub-panels (bar charts to the left and bottom of the main panel) show the number of isolates with each MIC (in line with the main panel). Yellow/blue coloring indicate which of these were predicted resistant/susceptible respectively, and black lines indicate cut-offs used to determine resistance classification (susceptible/resistant for EUCAST-based agar dilution, susceptible/intermediate/resistant for CLSI-based agar dilution).

Similarly, assessment of the individual contribution of other genetic features to the phenotype was challenging due to co-occurrence of features in the same isolate and the impact of some features on susceptibility varying both between isolates and within isolate repeats (Supplementary Figures S3, S4). For example, 4/9 isolates with ampC promoter mutations in the agar dilution subsample were both resistant and intermediate on repeat testing using CLSI-based agar dilution.

### WGS-derived resistance prediction in peri-breakpoint and non-peri-breakpoint isolates

As with routine AST, WGS predictions of reference-standard phenotypes were more accurate for non-peri-breakpoint MICs (EUCAST-based agar dilution: (≤4/2 mg/ml, ≥32/2 mg/ml), CLSI-based agar dilution: (≤4/2 mg/ml, ≥32/16 mg/ml)). For EUCAST-based agar dilution, WGS correctly identified resistance/susceptibility in 169/177 (95%) isolates with non-peri-breakpoint MICs, versus only 60/84 (71%) with peri-breakpoint MICs. Similarly, for CLSI-based agar dilution, excluding 79 intermediate isolates (16/8 mg/L), WGS correctly predicted 97/106 (92%) non-peri-breakpoint isolates, but predicted 43/76 (57%) isolates with MIC 8/4 mg/L as resistant.

Interestingly, however, there were three consistently resistant (EUCAST-based MIC ≥ 32/2mg/L, CLSI-based MIC ≥32/16 mg/L) and three consistently susceptible (EUCAST-based MIC ≤4/2mg/L, CLSI-based MIC ≤4/2 mg/L) discrepants. All three resistant discrepants were explained by complexities inferring phenotype from WGS. One had a novel *bla*_CTX-M_ variant (CTX-M-15-like, Ser130Gly mutation). Previous work on mechanisms of beta-lactamase inhibition suggests mutations at Ambler position(22) 130 likely lead to inhibitor resistance(7), and a similar mutation (Ser130Thr CTX-M-190) resulted in sulbactam and tazobactam resistance.(23) The other two isolates had antibiograms consistent with *ampC* hyper-production (cefoxitin resistant, ceftazidime resistant, cefepime susceptible), but we were unable to identify complete promoter sequences matching our reference (CP009072.1) in the region upstream of *ampC.* This may suggest insertion of alternative elements upstream of *ampC* could have led to both fragmented assemblies and have driven increased expression; however from WGS data alone it is difficult to distinguish if this has truly occurred or instead may be due to other undetected beta-lactamase resistance mechanisms. All three susceptible discrepants had beta-lactamases present at mildly elevated copy numbers (2.5-3.5x relative DNA coverage) leading to WGS prediction of resistance, which may be due to inherent unavoidable difficulty selecting cutoffs for predicting phenotype (Supplementary Figure S3B).

### Impact of individual resistance features on a continuous measure of susceptibility

Random-effects models were used to investigate the impact of test method and WGS-identified genetic elements on agar dilution log_2_ MICs simultaneously, and to create a WGS-based resistance prediction for comparison with phenotype (Supplementary Methods). Elements were categorised depending on frequency (Supplementary Table S5). The most predictive aspect of each element (including presence/absence of genes and/or promoter mutations and/or gene dosage) was selected using the Akaike Information Criterion (AIC) (Supplementary Methods). Interaction terms between genetic elements (reflecting saturation effects) and with test methodology (reflecting differential impact of the same genetic mechanism depending on the amoxicillin:clavulanate ratio) were included where p<0.05.

All beta-lactamases were associated with increased MICs in univariable models (Supplementary Table S6) and these associations generally persisted in multivariable models, although their magnitude decreased markedly (Figure 5, Supplementary Table S7). The largest effects of beta-lactamase presence/absence were for *bla*_OXA-1_ (a class 2d beta-lactamase, denoted blaOXA:2d) and members of the ‘other’ group of beta-lactamases, comprising either inhibitor-resistant beta-lactamases (N=10), or those with unknown impact on beta-lactam susceptibility (N=4) (Supplementary Table S5). These caused two-three fold and four-fold doubling dilution increases in EUCAST-based MIC respectively. The effects of non-inhibitor resistant *bla*_TEM_ (denoted *bla*TEM:2b) and *bla*_SHV_ (denoted blaSHV:2b) genes were more complex. For each, presence alone in an isolate was only associated with a small, often non-significant increase in MIC by either method (*bla*_TEM_: impact on change in log_2_(MIC) CLSI-based=+0.36 [p=0.01], EUCAST-based=+0.14 [p=0.51]; *bla*_SHV_: change in log_2_(MIC) CLSI-based=+0.03 [p=0.93], EUCAST-based=-0.61 [p=0.27]). However, for both, higher copy number (i.e. gene dosage) was associated with higher MIC. These effects were small but additive (e.g. EUCAST-based MIC change in log_2_(MIC) per doubling of copy number *bla*_TEM_=+0.79 [p<0.0001], *bla*_SHV_=+0.71 [p=0.004], detail in Supplementary Table S7A). Like beta-lactamases, all “significant” promoter mutations were associated with increased MICs (p<0.0001). In particular, “significant” *ampC* promoter mutations were independently associated with large increases in MIC (impact on change in log_2_(MIC) CLSI-based=+2.60 [p<0.0001], EUCAST-based=+4.25 [p<0.0001]). Interestingly, there was no clear change in MIC independently associated with suspected porin loss in our data (p≥0.06), despite porin loss being associated with a large effect in unadjusted analysis (Supplementary Table S6, change in log_2_(MIC) +2.28 (CLSI) and +4.17 (EUCAST)).

**Figure 5:**
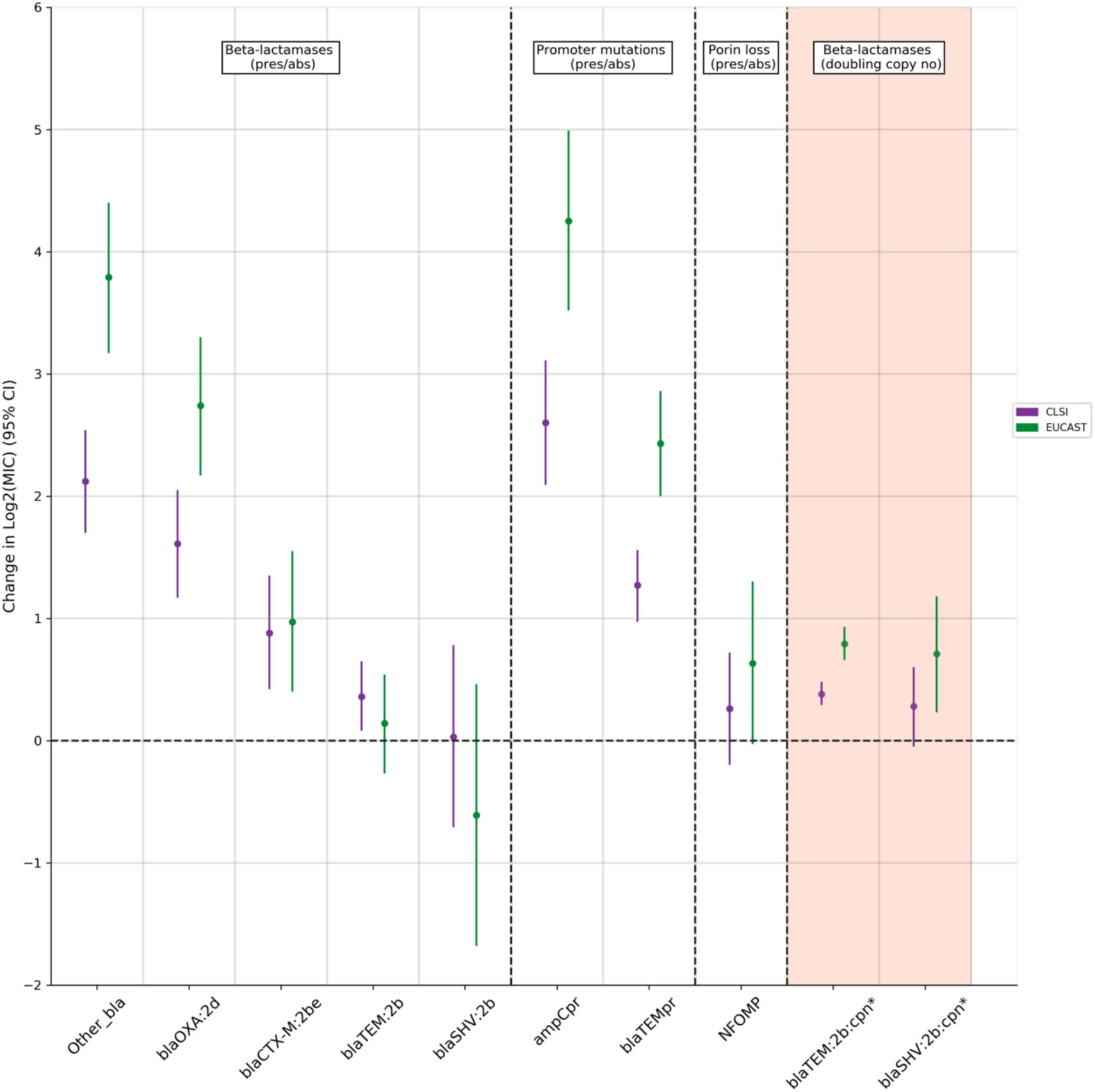
Changes in doubling dilution MIC independently associated with each feature/testing method (multivariable random-effects model). Note: Purple represents testing using 2:1 CLSI-based agar dilution (CLSI), and green using EUCAST-based agar diltuion. All elements except those denoted by * and shaded in orange are modelled as binary presence vs absence effects (see Supplementary Methods).), Other_bla (grouped other bla genes, includes bla_TEM-40_ (N=2), bla_TEM-30_(N=3), bla_CMY-2_ (N=3), bla_OXA-48_ (N=1), bla_TEM-190_ (N=1), bla_TEM-33_ (N=1), Supplementary Table 2) blaOXA:2d (Bush-Jacoby 2d, bla_OXA_), blaCTXM:2be (Bush-Jacoby 2be, CTXM), blaTEM:2b (Bush-Jacoby 2b, bla_TEM_), blaSHV:2b, (Bush-Jacoby 2b, SHV), ampCpr (ampC promoter mutation suggesting increased expression), blaTEMpr(bla_TEM_ hyper-producing promoter), NFOMP(non-functional ompF/ompC), blaTEM:2b:cpn (copy number) effect modelled as effect of doubling copy number, blaSHV:2b:cpn (copy number) effect modelled as effect of doubling copy number

Of note, when increased copy number effects were included, EUCAST-based testing methodology accentuated increases in MIC caused by genetic resistance features other than for suspected porin loss and presence of *bla*_CTX-M_ genes (the blaCTX-M:2be group)(p_heterogeneity_≤0.05). EUCAST-based methodology however was also associated with increased between and within sample standard deviation (Supplementary Table S7B).

### Predictions of MIC in an independent validation set

Final EUCAST-based agar-dilution model estimates were then used to predict MICs for the 715/976 non-subsample isolates, which were then compared with BD Phoenix MICs. MIC predictions were in agreement for 557/715 (78%) isolates and in essential agreement (within ±1 doubling dilution) for 691/715 (97%). However, these 715 non-subsample isolates included 11isolates which contained resistance mechanisms not present among the agar-dilution subsample isolates from which the model was derived (e.g. different beta-lactamase variants). Excluding these, prediction performance was similar, with agreement for 554/704 (79%) isolates (Figure 6) and essential agreement for 683/704 (97%) isolates. Similarly to comparisons between the different antimicrobial susceptibility testing methods (Figure 3), agreement between predicted and observed resistant/susceptible classifications was lower (90%) despite having high essential agreement of MICs. While overall performance was good, three isolates had predicted MICs three doubling dilutions lower than observed. One had an unusual yet reproducible phenotype [ampicillin susceptible, amoxicillin-clavulanate resistant]. This rare phenotype has generally been found in non-*E. coli* Enterobacteriaceae, and is thought to be due to either mechanisms of *ampC* induction(24) or the differential activity of amoxicillin and ampicillin.(25) We were unable to identify a clear causative mechanism in this isolate; however, of note, it was the only isolate to contain a -11 C>T *ampC* promoter mutation.. The other two both had observed MIC ≥32/2mg/L but only contained a low copy number *bla*_TEM-1_ and had predicted MIC 8/2mg/L.

**Figure 6.**
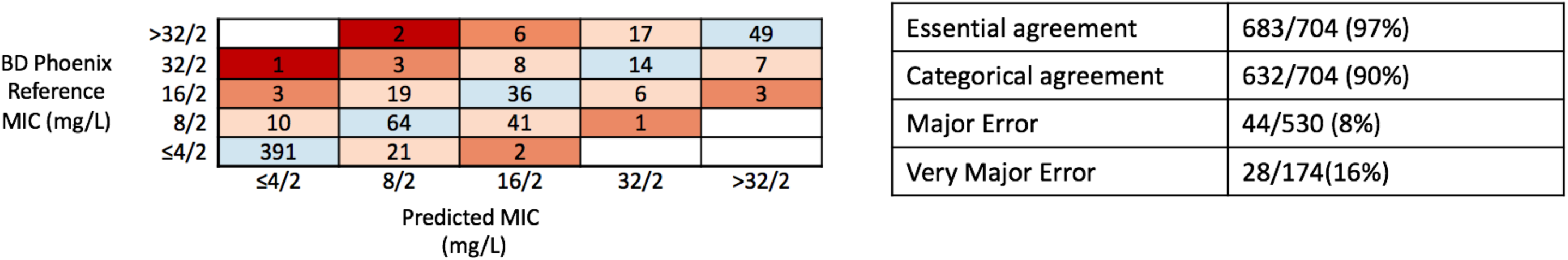
Model based MIC prediction for non-subsample isolates (N=704) Note: blue shading indicates correct correctly predicted observed AST MIC (554/704 (79%) isolates), light pink predicted within one doubling dilution (total 683/704 (97%) isolates, essential agreement), orange within two doubling dilutions (total 701/704 (100%)) and red greater than 2 doubling dilutions. Excluding 11 isolates with resistance mechanisms not included in the agar-dilution subsample on which the prediction model was derived (similar overall performance including these).

## Discussion

Decisions about broadening recommended empiric antimicrobial regimens from amoxicillin-clavulanate are currently being made based on unclear AST data which appears poorly concordant with WGS-identified determinants of beta-lactamase resistance. Here, we have demonstrated that this lack of concordance is not due to unknown genetic features or inherent phenotyping problems as previously hypothesized.(26) Instead, it appears to arise from poor interpretation of how known genetic mechanisms of resistance impact phenotype. Contrary to the often assumed paradigm that beta-lactam resistance is generally due to the presence/absence of specific beta-lactamases alone, mechanisms of resistance to amoxicillin-clavulanate seen regularly in a large unselected clinical dataset were multifactorial, resulting from combinations of multi-copy beta-lactamase genes, mutations in resistance gene-associated promoters, and inhibitor resistance mechanisms. The individual effects of some of these features on MIC were small, variable and additive, resulting in only minor shifts around clinical breakpoints. This potentially explains inconsistencies on repeated phenotyping, and may be a consequence of the genetic basis of resistance rather than an inherent test weakness. A further corollary is that discrepancies between genotypic predictions and phenotype are inevitable when using susceptible/resistant binary classifications. Finally, the phenotypic testing methodology significantly affected the magnitude of the effect of these resistance features on the MIC. These issues, when combined, resulted in inconsistent binary phenotypes despite reliable MICs, and consequently led to inevitable suboptimal concordance both between different phenotypic testing methodologies and also with WGS-based susceptibility/resistance predictions An alternative approach would be to use WGS to predict MICs directly. We demonstrated this was possible by predicting the MIC to within one doubling dilution (essential agreement) of the observed MIC for 97% of isolates from a population-representative set of *E. coli* BSI.

Our study highlights the importance of isolate sampling frame, phenotyping method and breakpoint selection. A previous study of 76 *E. coli* isolated from cattle (19), which reported high sensitivity and specificity of WGS to predict amoxicillin-clavulanate resistance, contained highly-resistant isolates (30% containing bla_CMY-2_), and only attempted to predict CLSI-defined resistance (>32/16 mg/L). In contrast, in our study, similar to other population representative studies of human isolates,(6, 14) only a small proportion of amoxicillin-clavulanate resistance was due to inhibitor-resistant beta-lactamases, with most of the resistance being due to hyper-production of beta-lactamases. Further, while EUCAST argue that pharmacodynamic data support choice of breakpoint and clavulanate concentration (9, 27), there is no definitive evidence as to which method has stronger associations with either clinical outcome or genotype. We therefore assessed WGS against both commonly used methods (EUCAST, CLSI).

Compared with other studies of BL/BLIs and *E. coli* causing human infections, we found less BL/BLI resistance was accounted for by inhibitor-resistant beta-lactamases.(20) To identify resistance in our population-representative set of isolates, we found it critical to consider genetic features that alter expression of beta-lactamases. Although the individual effects of some of these features on MICs were small, they were important, because MICs for many isolates were close to the breakpoint. Further, given the small size of these effects and effects of testing methodology, isolates could exhibit either susceptible or resistant phenotypes on repeat testing, supporting the concept of an “intermediate” phenotype, which is not accounted for in the EUCAST guidelines. The discrepancies between EUCAST and CLSI phenotypes we observed were similar to previous studies,(3) suggesting that phenotypic interpretation for one of our most commonly used clinical antibiotics remains open to question. The one mechanism for which we found no evidence of effect was porin loss: this may reflect difficulty detecting these effects from WGS alone or be a simple power issue given rarity of this mechanism in our population-based sample, since porin loss has been associated with raised MICs to other BL/BLI combination antibiotics in isolates containing *Klebsiella pneumoniae* carbapenemases.(28)

The main study strengths are the large, population-representative sampling frame; detailed, replicated, reference-grade phenotyping for a substantial subset of isolates on which prediction models were developed (“training set”) and a large number of additional isolates with single phenotypes assessed by a commercial clinically accredited platform (“test set”); detailed and complete genotyping; and the statistical modelling. While model performance on other datasets with geographical differences in resistance mechanisms and prevalence is unknown, its good performance in unselected clinical isolates with resistance mechanisms commonly seen in practice (in contrast to many previous studies of WGS-based resistance prediction(29)) suggest it may be generalizable. Further, the consistency of individual findings with previous literature (including inconsistent phenotype(30), mechanisms of genetic resistance(31) and poor performance of beta-lactamase presence/absence-only prediction for BL/BLI resistance (20, 32)) provide confidence that the combined results may apply across different settings. A further study strength was our use of more complete representations of the mechanisms of beta-lactam resistance. Compared to other studies of WGS-based resistance prediction which have either just used presence/absence of beta-lactamases or machine learning methods directly on sequence data, our models more easily align with traditional approaches of studying antimicrobial resistance and provide interpretable estimates of the direct effects of different mechanisms. A limitation is that we could only investigate proxies for some important genetic features, e.g. increased DNA copy number leading to increased expression. WGS is unable to directly quantify these effects, which thus require additional characterization by alternative methods, leading to concern that resistance prediction from WGS alone would be highly challenging. In practice however, the good predictive performance of our model using relatively simple proxies suggests many of these features can indeed be approximated from WGS data. Modelling associations between resistance features and MIC directly allowed us to avoid inferring the phenotype from the genotype using pre-specified rules and account for the effects of multiple features existing in individual isolates. The complexity of the underlying associations we discovered highlights the challenges facing standardized methods for predicting resistance across multiple drugs and species,(33) and the need for automated approaches based on machine learning to take into account proxies for increased expression.

The main limitations of this study relate to its size. While we determined repeat agar dilution phenotypes for a relatively large number of isolates (n=261) compared to other studies,(30) many resistance elements were still rare. This had three important consequences: some infrequent features had to be categorized together for modelling, interactions between all combinations features (e.g. combinations of beta-lactamases) could not be definitively assessed, and some mechanisms present in our testing set were not present in the training set (e.g. some rare known beta-lactamase variants, many only in a single isolate, Supplementary Table S2) and so their effect could not be estimated. Model results however suggested these had limited consequences. Firstly, the features causing the greatest MIC increases were those traditionally associated with amoxicillin-clavulanate resistance,(7) their specific impact being modelled here for the first time. Secondly, only a small number of isolates had resistance mechanisms not seen in the training dataset (N=11), meaning impact on performance was minimal (essentially their effect was assumed to be 0). This issue is inevitable given the substantial diversity of *E. coli* and incomplete knowledge of resistance mechanisms, but the excellent performance in the remaining 704/715 isolates suggests the vast majority of clinical isolates could be amenable to WGS-based MIC prediction, leaving a much smaller, more tractable number of isolates needing additional phenotypic investigation. Another potential limitation was the use of agar dilution as our reference-standard phenotype, a method which, while previously endorsed by EUCAST,(34) is no longer recommended, with broth microdilution now recommended instead. By contrast CLSI still considers agar dilution as equivalent to broth microdilution.(35) Reassuringly, differences we found between the BD Phoenix and agar-dilution were similar to a previous study comparing BD Phoenix with reference-standard broth microdilution,(36) suggesting this would have relatively little impact on our overall results.

In summary, amoxicillin-clavulanate resistance in *E. coli* is quantitative, rather than qualitative; in reality, resistance is a continuum built up by many individual features inevitably resulting in poor reproducibility and suboptimal concordance with binary classifications. WGS can identify the causes of amoxicillin-clavulanate resistance in *E. coli* provided the approach is extended to consider the complicated, polygenic, and expression-related nature of this resistance. This suggests a genetic approach could offer a less assay dependent way to assess amoxicillin-clavulanate resistance. With renewed interest in using BL/BLIs to treat highly drug-resistant infections, our study has implications for both clinical practice and research. Given susceptibility phenotypes are highly dependent on the phenotypic method used, they must be interpreted with caution. Genetic approaches have the potential to circumvent this issue. Importantly however, the assumption that BL/BLI resistance is binary (susceptible/resistant) may be unhelpful as the same underlying resistance feature can be associated with MICs just below or just above the breakpoint. Given the variability and complexity in both the underlying mechanisms and resulting phenotype, a more transparent approach considering background genetic features, expression levels of beta-lactamases, MIC values and clinical syndrome, is likely needed to guide management decisions.

## Materials and Methods

### Study population and routine microbiological processing

*E. coli* isolated from all monomicrobial or polymicrobial blood cultures at Oxford University Hospitals (OUH) NHS Foundation Trust between 01/Jan/2013-31/Aug/2015 were included, excluding repeat positive cultures within 90-days of an index positive. Automated AST was performed in the routine laboratory (BD Phoenix; Beckton, Dickinson and Company) and MICs interpreted using EUCAST breakpoints. Data were extracted from the Infectious Diseases in Oxfordshire Research Database (IORD)(37) which has Research Ethics Committee and Health Research Authority approvals (14/SC/1069, ECC5-017(A)/2009).

### DNA extraction and sequencing

Isolates were re-cultured from frozen stocks stored in nutrient broth plus 10% glycerol at -80°C. DNA was extracted using the QuickGene DNA Tissue Kit S (Kurabo Industries, Japan) as per manufacturer’s instructions, with an additional mechanical lysis step (FastPrep, MP Biomedicals, USA) immediately following chemical lysis. A combination of standard Illumina and in-house protocols were used to produce multiplexed paired-end libraries which were sequenced on the Illumina HiSeq 2500, generating 151bp paired-end reads. High quality sequences (Supplementary Methods) were de-novo assembled using Velvet(38) as previously described.(39) *In silico* Achtman(40) multi-locus sequence types (MLST) types were defined using ARIBA.(21)

### Evaluating the importance of genetic features that modify effective beta-lactamase concentration

We identified components of two genetic resistance prediction algorithms for amoxicillin and amoxicillin-clavulanate (Table 2, Supplementary Methods) using ARIBA(21) (default parameters) and tBLASTn/BLASTn.(41) The “basic” prediction used only presence/absence of relevant genes in the Resfinder(17) database, and the “extended” prediction additionally included *bla*_TEM_ and *ampC* promoter mutations, estimates of DNA copy number and predicted porin loss-of-function. High DNA copy number was used as an indicator of possible gene duplication or high plasmid copy number, which are both known to cause increased beta-lactam resistance.(42, 43) We made no attempt to distinguish between these two causes due to the limitations of short-read sequencing data. For *bla*_TEM_ and *ampC* promoters, sequences identified using ARIBA/BLASTn were searched for variant sites and regions previously associated with significantly increased expression.(44–46) For transmissible resistance genes, we estimated DNA copy number by comparing mapping coverage with the mean coverage of MLST genes and defined a relative coverage of >2.5 as increased copy number (based on receiver-operator-curve (ROC) analysis, Supplementary Methods; Supplementary Figure S3A). Finally, sequences found by ARIBA using reference *ompC* and *ompF* sequences (RefSeq: NC_000913.3) were inspected for features such as indels and truncations suggesting functional porin loss.

**Table 2:** Resistance prediction feature performance

| Feature | Prediction | Number of isolates resistant by routine AST/total isolates with this feature (%) |  |
| --- | --- | --- | --- |
|  |  | PPV | PPV when in isolation<sup>+</sup> |
| **Beta-lactamases** |  |  |  |
| **1** Any class C or D serine beta-lactamase | Basic | 66/69 (96%) | 32/34 (94%) |
| **2** Any inhibitor resistant class A beta-lactamase | Basic | 12/12 (100%) | 2/2 (100%) |
| **Promoter mutations** |  |  |  |
| **3** Non-P3 *bla*<sub>TEM</sub> promoter associated with *bla*<sub>TEM</sub> hyper-production | Extended | 48/49 (98%) | 29/30 (97%) |
| **4** - *ampC* promoter mutation associated with *ampC* hyper-production | Extended | 20/20 (100%) | 13/13 (100%) |
| **Increased DNA copy number** |  |  |  |
| **5** Relative coverage of any transmissible beta-lactamase > 2.5* | Extended | 184/227 (81%) | 128/167 (77%) |
| **Decreased permeability<sup>o</sup>** |  |  |  |
| **6** Features suggesting disruption of either *ompC* or *ompF* in an isolate containing an additional beta-lactamase (see supplementary methods) | Extended | 19/22 (86%) | 2/2 (100%) |

+: PPV restricted to isolates not predicted as amoxicillin-clavulanate resistant by any other feature
\*: Cut-off chosen following a receiver operating curve (ROC) analysis (Supplementary Methods, Supplementary Figure 3A)
<sup>o</sup>: Beta-lactam resistance features in isolates shown in Supplementary Tables S2, S3, S4
Note: PPV=positive predictive value

### Evaluating the impact of different phenotypic methods

A subset of 291 isolates were selected for replicate agar dilution phenotyping using random sampling within strata defined by phenotype-genotype combinations (see Supplementary Methods for full sub-sampling procedure, Supplementary Figure S1A,B). Replicate agar dilution phenotyping used clavulanate concentration and MIC interpretation according to both EUCAST and CLSI guidelines. The aim was to explore reasons for discordance between observed phenotype and predictions made on the basis of beta-lactamase gene presence/absence alone. The subsampling therefore aimed to enrich for several groups of isolates, including resistant (by BD Phoenix) isolates both with and without beta-lactamases identified from WGS, isolates with peri-breakpoint MICs and susceptible isolates containing beta-lactamases. For each method, sub-cultures (from frozen stocks) were tested in triplicate using ISO-Sensitest agar plates containing amoxicillin and clavulanate in a 2:1 ratio (CLSI) or a fixed concentration of clavulanate (2 mg/L) (EUCAST), with *E. coli* controls ATCC25922 (wild type) and ATCC35218 (TEM-1 beta-lactamase producer).(47) For additional quality control, bacterial isolates were plated on sheep blood agar and incubated overnight at 37°C to check purity, with isolates excluded if multiple colonial morphologies were seen. Isolates were included in analyses if two or more MICs for each of amoxicillin, EUCAST-based amoxicillin-clavulanate and CLSI-based amoxicillin-clavulanate were in essential agreement (i.e. a minimum of 2/3 for each drug; see Supplementary Methods). Isolates with less than two MICs for each of the tests passing quality control were tested an additional time to reduce the risk of selection bias against isolates with genetic mechanisms causing variable expression and underestimating natural phenotypic variability. For each included isolate, susceptibility classification for that isolate was defined using the “upper median” MIC (choosing the higher MIC when the median lay between two MIC readings) of the test repeats.

### Modelling and predicting MICs

Random-effects models (Stata 14.2; StataCorp LP, 2015) were used to investigate the impact of test method and WGS-identified genetic elements on agar dilution log_2_ MICs simultaneously (additional details in Supplementary Methods). Elements were categorised depending on frequency (Supplementary Table S4). Models included method-specific random-effects for each isolate and testing batch, and method-specific (heteroskedastic) errors. All genetic element categories were included *a priori*, but the most predictive effects of each (including presence/absence of genes and/or promoter mutations and/or gene dosage) were selected using the Akaike Information Criterion (AIC) (Supplementary Methods). Lastly, interaction terms between genetic elements (reflecting saturation effects) and with test methodology (reflecting differential impact of the same genetic mechanism depending on the amoxicillin:clavulanate ratio) were included where p<0.05. Final estimates were then used to predict MICs in all non-subsample isolates and in non-subsample isolates which did not contain resistance features not present in the agar dilution subsample. Predicted MICs were then compared to routine laboratory phenotypes from the BD Phoenix.

## Acknowledgments

This work uses data provided by patients and collected by the NHS as part of their care and support. We thank all the people of Oxfordshire who contribute to the Infections in Oxfordshire Research Database. Research Database Team: R Alstead, C Bunch, DCW Crook, J Davies, J Finney, J Gearing (community), L O’Connor, TEA Peto (PI), TP Quan, J Robinson (community), B Shine, AS Walker, D Waller, D Wyllie. Patient and Public Panel: G Blower, C Mancey, P McLoughlin, B Nichols. We would also like to thank the HPRU Steering Group (N French, C Marwick, J Coia, M Sharland).

## Author contributions

TJD, NS, MJE, NW, DWC, TEAP, MFA and ASW designed the study. KJ, MA(OUH), MM, TPQ obtained the automated susceptibility phenotypes from archived BD phoenix records. TJD, and MA(APHA) performed agar dilution on samples. DG and AV sequenced isolates. TJD, HP and JS ran resistance genotype prediction on samples. TJD, AS, NS, OE, RB and AM interpreted the genetic results and established rules regarding the relationship with phenotype. TJD and ASW fitted random-effects models to the data. TJD, NS, AS, PF, ASW and MFA prepared the first draft. All authors commented on the data and its interpretation, revised the content critically and approved the final version.

## Data and materials availability

Sequences used in the study are made available at PRJNA540750. MIC data and code used for this analysis are available at https://github.com/TimothyJDavies/reconciling_the_potentially_irreconcilable

## List of contents

Supplementary Materials and Methods

Supplementary Figure S1: Sampling frame and sample selection

> A: Sampling frame and included isolates

> B: Agar dilution subsample selection

Supplement Figure S2: Isolate STs, resistance mechanisms and amoxicillin-clavulanate phenotyping for:

> A: The full dataset (N=976)

> B: The agar dilution subsample (N=261)

Supplementary Figure S3: Estimating the coverage copy number cut-off

> A: ROC curve for selection of DNA copy number threshold for the extended resistance prediction

> B: Association between DNA copy number and agar-dilution phenotype in isolates containing only non-inhibitor resistant beta-lactamases

> C: Persistent classification errors even when choosing optimal coverage cut-offs due to normal distribution of MICs, demonstrated in isolates containing only low copy number bla_TEM-1_

Supplementary Figure S4: Maximum MIC doubling dilution difference across repeats by method for subsample isolates

Supplementary Table S1: Beta-lactam antibiograms

Supplementary Table S2: Beta-lactamases

Supplementary Table S3: Promoter sequences

> A: *ampC* promotors

> B: amoxicillin-clavulanate MICs of *ampC* hyper-producing isolates

> C: *bla*_TEM_ promotors

Supplementary Table S4: Disrupted porin genes

Supplementary Table S5: Random-effects model categories

Supplementary Table S6: Unadjusted resistance feature effects

Supplementary Table S7: Random-effects model results

> A: Independent predictors of agar dilution log2 MIC

> B: Estimated variation in MIC according to different sources from the random-effects model

