## Supplementary Materials for "Reconciling the potentially irreconcilable? Genotypic and phenotypic amoxicillin-clavulanate resistance in *Escherichia coli*"

**Supplementary Methods**

**Included samples and subsampling procedure**

*Sampling frame and included isolates*

We attempted to include at least one *E. coli* isolate from every case of *E. coli* bloodstream infection (BSI) (excluding repeat isolations within 90 days of an index positive culture) at Oxford University Hospitals (OUH) NHS Foundation Trust between 01/Jan/2013-31/Aug/2015. Over the study period, there were 1039 distinct *E. coli* BSI episodes, from which 1054 *E. coli* were isolated. We were able to sequence and had automated antimicrobial susceptibility testing (AST) data for 1008/1054 (96%) isolates, representing 1000/1039 (96%) cases of BSI (Supplementary Figure S1A). MIC data was used to infer phenotype using EUCAST breakpoints (version 8.1).[1] Of these, 723 isolates (01/Jan/2013 – 31/Dec/2014) had complete sequencing data and automated AST phenotypes available at the time of selection for the agar-dilution subsample. Overall for the main study, post sequencing quality control we included 976/1054 (93%) isolates representing 968 of the possible 1039 (93%) *E. coli* BSI.

*Agar dilution MIC sub-study sample selection*

At the time of subsample selection, an initial WGS resistance prediction was generated using BLASTN[2] searches on *de-novo* velvet assemblies[3]. The search used an extended version of the database used by Stoesser *et a*l[4], searching for inhibitor resistant beta-lactamases (basic prediction) and *bla*_TEM_ promoter mutations. Of note, this classification did not include *ampC* promoter mutations, copy number and porin functionality. Using a combination of this initial prediction, whether the sample contained a beta-lactamase and the initial laboratory phenotype, samples were classified into 9 strata for subsampling (Supplementary Fig. S1B). Samples were then selected at random within each strata, but enriching for several phenotype-genotype combinations. Of note, group 9 represented all piperacillin-tazobactam resistant samples which were not selected as part of another group.

**WGS resistance prediction algorithms**

*Method*

Sequencing data for each isolate was interrogated using ARIBA[5] (using default settings) with an extended version of the Resfinder database[6] (base database accessed 16^th^ November, 2017) which additionally included a template *ampC* promoter sequence, *bla*_TEM_ promoter sequence and porins *ompF* and *ompC*. Features were deemed present if

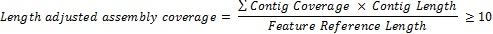

Information on heterozygous hits and disruption ARIBA assemblies were not used in any resistance prediction outside of porin genes (so as not to require complete assemblies/consensus sequences for genes found, a known issue when looking for resistance features in gram negative organisms[7]). However, these were fully investigated on discrepancy checking, including analysing the resistance profile of all predicted heterozygous alleles present in each isolate.

*Basic algorithm: Inhibitor resistant beta-lactamases*

Analogous to Stoesser *et al*[4], isolates were deemed inhibitor (clavulanic acid) resistant if they contained one of the following:

- Any ambler class C or D beta-lactamase gene (e.g. *bla*_OXA-1_, *bla*_CMY-2_)
- Any inhibitor resistant ambler class A beta-lactamase (e.g. *bla*_TEM-30,_ *bla* _SHV-10_ *)*

*Extended algorithm: bla_TEM_ promoter mutations.*

To identify potential *bla*_TEM_ promoter sequences associated with increased *bla*_TEM_ expression, we searched each isolate’s sequence data for sequences similar to the Lartigue *bla*_TEM_ P3 promoter.[8] Sequences found were then compared to each of the Lartigue promoter types, and the closest match was selected. Resistance was inferred if the closest match was any non P3 promoter.

*Extended algorithm: ampC promoter mutations.*

Similarly to the *bla*_TEM_ promoter, we searched each isolate’s sequence data for sequences similar to the *ampC* promoter present in ATCC 25922. Sequences found were inspected for variants known to increase chromosomal *ampC* expression in *E. coli.*[9] Isolates with any of these variants were predicted resistant by the extended algorithm. In addition, as *ampC* and its promoter should be universally present among *E. coli*, we noted where we were unable to find *ampC* promoter sequences for discrepancy analysis.

*Extended algorithm: DNA copy number*

*Generating the metric*

For all transmissible genes, we generated a DNA copy number metric from ARIBA output, defined as

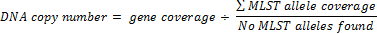

where gene coverage was defined as the coverage of the longest contig in the ARIBA assembly. For example, a DNA copy number of 1 suggests the gene is present in the same quantities as the MLST genes (i.e. 1 per cell) and a DNA copy number greater than 1 suggests the gene is present in higher quantities than the MLST genes (i.e. > 1 per cell).

*Reliability of the DNA copy number metric*

Measurement error in the copy number was estimated to be small (standard deviation = 0.17 (95% CI 0.14, 0.21)) based on copy numbers estimated for 46 elements from 9 isolates sequenced in duplicate as part of quality control. Therefore the absolute estimated DNA copy number was used in subsequent analyses.

*Choice of DNA copy number threshold for the extended resistance prediction*

This cut-off was chosen based on a Receiver Operating Characteristic (ROC) analysis of all 328 isolates containing a single beta-lactamase as the only potential cause of resistance (Supplementary Figure 3A). Maximal Youden index, maximal Liu index and minimal distance to (0,1) all selected cut-offs between 2.3 and 2.4 (bootstrap 95% confidence intervals contained within 1.7-3.0). We rounded to the nearest half number for easier interpretation, defining 2.5 as the threshold. The fact that the threshold is not an integer suggests the gene may be present in varying copies in different cells (e.g. some cells containing 2, and others containing 3).

*Extended algorithm: porin loss of function*

To investigate potential loss of function of porin mutations, we searched each isolate’s sequence data for sequences similar to reference *ompC* and *ompF* sequences (RefSeq: NC_000913.3). Given these sequences should be ubiquitous in *E. coli*, we defined being unable to find and assemble a complete coding sequence for either as likely signifying porin loss. The presence of any of the following factors were used to determine this

- Absence of any of “unique contig”, “complete gene found” or “assembled into one contig” ARIBA flags
- Presence of any of “hit both strands: “region assembled twice”, “scaffold graph bad” ARIBA flags
- If length of the found sequence != length of the reference sequence
- If percentage identity of protein alignment to reference sequence < 90%
- If the found sequence contains a mutation causing truncation (i.e. the entire found sequence must be coding)

Given reduced permeability is generally thought to contribute to multi-mechanism resistance in isolates[10], suspected porin loss was only used to predict amoxicillin-clavulanate resistance when the isolate additionally contained a beta-lactamase.

**Sequencing quality control**

WGS data quality was ensured by obtaining and assessing the following metrics from the data.

- Sequencing data for each isolate needed to contain > 1000000 reads
- The top species (as identified using Kraken[11])
- The *de-novo* assembly had to meet the following criteria
  - total length had to be between 4 and 6 megabases long
  - the N50 (minimum contig length needed to cover 50% of the genome) was greater than 50000
  - the assembly contained less than 250 contigs greater than 500bp long
  - the assembly contained less than 800 contigs
- The mapping of WGS data to to *E. coli* CFT073 (AE014075.1) had to meet the following criteria
  - > 60 % of reads mapped to reference
  - > 60 % of reference bases called as “A”, “T”, “C” of “G”
- There were no signs of mixture identified by in-silico MLST typing by ARIBA
  - No heterozygosity of MLST alleles for MLST alleles that were present at for 15x coverage

**Agar dilution MIC quality control**

Agar dilution MIC testing for each isolate selected to be part of the agar dilution MIC subsample was performed following British Society for Antimicrobial Chemotherapy (BSAC) guidelines, which were widely in use in the UK during the study.[12] For additional quality control, phenotyping was attempted a minimum of 3 times for each isolate. To be deemed a satisfactory result, a test had to fulfil the following criteria:

- plate control ATCC25922 had amoxicillin MIC in range 2-8 mg/l;
- plate control ATCC25922 had 2:1 ratio amoxicillin/clavulanate MIC 2/1-8/4 mg/l ;
- plate control ATCC25922 had fixed 2mg/l clavulanate MIC 2/2-8/2 mg/l;
- plate control ATCC35218 had 2:1 ratio amoxicillin/clavulanate MIC 4/2-16/8 mg/l;
- plate control ATCC35218 had fixed 2mg/l clavulanate MIC 4/2-32/2 mg/l;
- only 1 visible colony type seen on purity plate (sheep’s blood agar).

Then for each isolate, it was only included in the agar dilution subsample if we obtained 2 or more MICs in essential agreement for all of amoxicillin, fixed-based amoxicillin-clavulanate and ratio-based amoxicillin-clavulanate. Any test with less than two “passing” sets of agar dilution results were re-tested (for all of amoxicillin, fixed and ratio tests) and included if it then had two or more MICs in essential agreement for all of amoxicillin, fixed-based amoxicillin-clavulanate and ratio-based amoxicillin-clavulanate. The reasoning behind this protocol of repeats and quality control requirements was to obtain MICs with at least the support of a duplicate test (suggesting that the sample was not mixed), but also to avoid creating selection bias by dropping samples with greater variability in MIC since this could select against some mechanisms of resistance with variable expression. Of note while the phenotype was repeated a minimum of three times, our quality control and retesting procedure resulted in some isolates having even numbers of tests for one or more antibiotics, necessitating the use of the “upper median MIC” (choosing the higher MIC when the median lay between two MIC readings) as described in the main text.

**Random-effects models**

*Categorization of resistance mechanisms identified from WGS for modelling*

Supplementary Tables S2 and S3 detail the beta-lactamase genes (Supplementary Table S2), *ampC* promotors (Supplementary Table S3) and *bla*_TEM_ promotors (Supplementary Table S3) found in the study isolates. In a subset of isolates undergoing agar dilution MICs, we estimated the individual effect of these resistance mechanisms. Some mechanisms occurred in very few isolates in this agar dilution subsample, and so could not be considered individually. Beta-lactamases found as “complete genes” by ARIBA were categorised by family and Bush-Jacoby[13] classification (Supplementary Table S5). *bla* genes in categories with fewer than 10 isolates and non-complete *bla* genes present in the agar dilution subsample were reclassified as an “other” group. This resulted in 5 categories of beta-lactamases

- blaTEM:2b containing the class 2b *bla*_TEM_ genes
- blaCTXM:2be containing the class 2be *bla*_CTX-M_ genes
- blaOXA:2d containing *bla*_OXA-1_ genes (a class 2d enzyme)
- blaSHV:2b containing *bla*_SHV-11_ genes (a class 2b enzyme)
- Other_bla: containing the “other” beta-lactamases. In the agar dilution subsample this group contained 10 inhibitor resistant beta-lactamases, and 4 beta-lactamases with unknown impact on amoxicillin-clavulanate resistance.

Given the large number of individually relatively rare *ampC* promoters, they were grouped and classified as “ampCpr*”,* equalling 1 if containing a resistance conferring mutation, 0 if otherwise. Likewise, *bla*_TEM_ promoters were grouped and classified similarly as “blaTEMpr”. Isolates with both a suspected loss of function of either *ompC* or *ompF* (see above) and a beta-lactamase were classified as having a significant permeability effect (NFOMP(non-functioning omp)=1, 0 otherwise).

*Model selection: variance components*

Random-effects models were constructed to relate each MIC (modelled on the log_2_ scale, so a 1 unit increase represents a doubling) to various genetic elements and were fitted using maximum likelihood estimation. Each test result was included as an observation, accounting for repeated tests of the same isolate using isolate-level random effects. Left censoring of co-amoxiclav MICs at 2/2 (EUCAST) and 2/1 (CLSI) mg/L, and right censoring at >128/2 mg/L (EUCAST only) in a small proportion of isolates was dealt with by dividing or multiplying the observed lower limit by half respectively (i.e. subtracting or adding 1 on the log2 scale[14]). Sets of agar plates containing varying concentrations of amoxicillin:clavulanate (fixed clavulanate 2mg/L (EUCAST) and 2:1 ratio clavulanate (CLSI)) were prepared in batches, which were then used to test 55 isolates for each batch. One source of variation in MICs is therefore the test batch. Another source of variation is the isolate itself. As not all batches contained the same isolates due to repeat testing, isolates were not nested within the same batch. The random effects structure was therefore chosen by taking the variance components structure with the lowest Akaike Information criterion (AIC) from random effects for isolate only, batch only, separate and crossed random effects for isolate and batch. The best fit was cross classified, with each individual MIC nested within both isolate and batch, and neither of isolate/batch nested in the other. Including the main 3 MLSTs individually or together as fixed effects did not improve model fit (p>0.14) and so MLSTs were not considered further.

*Model selection: main fixed components*

Given the need to control for the effects of the different clavulanate ratios intrinsic to the two test methods (EUCAST vs CLSI), models a-priori included a main fixed effect for test method and each genetic element, initially including all genetic elements as binary presence/absence (classical interpretation; multivariable model). We then considered whether there was evidence for additional effects of gene dosage. Modelling considered each beta-lactamase in turn, with the most common first (i.e. *bla*_TEM_, then *bla*_CTX-M_, *bla*_OXA_, *bla*_SHV_), for each of the following:

- Binary (presence/absence, classical interpretation) (default)
- Binary (presence/absence) + linear effect of increasing copy number when present (each absolute unit higher copy number has the same effect on log2 MIC)
- Binary (presence absence) + log2-linear effect of increasing copy number when present (each doubling of copy number has the same effect on log2 MIC)

The best fitting model for each beta-lactamase gene was chosen based on the AIC. To reduce the influence of outliers, we truncated one extreme copy number (54.8) for SHV2b (in rest of isolates median 1.77, IQR (1.70,9.05)) to the 95^th^ centile (16.27). There were no other clear copy number outliers.

Following this analysis, beta-Lactamase categories were modelled as follows

- blaTEM:2b – Binary (presence/absence) + Log2(copy number for copy number > 1)
- blaCTXM:2be – Binary(presence/absence)
- blaOXA:2d – Binary(presence/absence)
- blaSHV:2b – Binary (presence/absence) + Log2(copy number for copy number > 1)
- Other_bla – Binary (presence/absence)
- blaTEMpr – Binary (presence/absence of significant mutations )
- ampCpr – Binary (presence/absence of significant mutations )
- NFOMP – Binary (presence/absence of evidence of disruption)

*Model selection: interactions*

As defined, the blaTEMpr and NFOMP (porin loss) factors already represented an interaction with other beta-lactamases, since by definition they can only have an impact in the presence of a beta-lactamase; therefore these were omitted from further investigation of interactions. Given the importance of test method, we included all first order interactions between test method and all other main genetic factors in the model. All individually were p<0.05 on a likelihood ratio test comparing multivariable models with and without each specific interaction, supporting this choice, with the exception of the interaction between test method and NFOMP or blaCTX-M:2be which may be due to the smaller number of isolates with these mechanisms (Supplementary Table S5). Other first order interactions between included genetic elements were assessed if they were found together in 5 or more isolates, and were selected by forward selection using likelihood ratio tests with p<0.05. These included interactions all reduced MIC when multiple elements were found together, reflecting saturation effects whereby the combined effect of having two genetic elements which individually both increase MIC is not the additive effect of both, but slightly less than this (see equation below).

Finally, we considered whether the random effects (by batch and isolate) varied according to test methodology. Including heteroskedastic residual errors (i.e. allowing the residual error of each test MIC to vary across test methodology), and allowing the random error associated with each isolate’s MIC and the random error associated with batch effects to vary according to test method improved model AIC and so were included, resulting in a final model as shown below. Covariance between test method was only estimable on isolate level random errors due to small numbers of clusters at batch level.

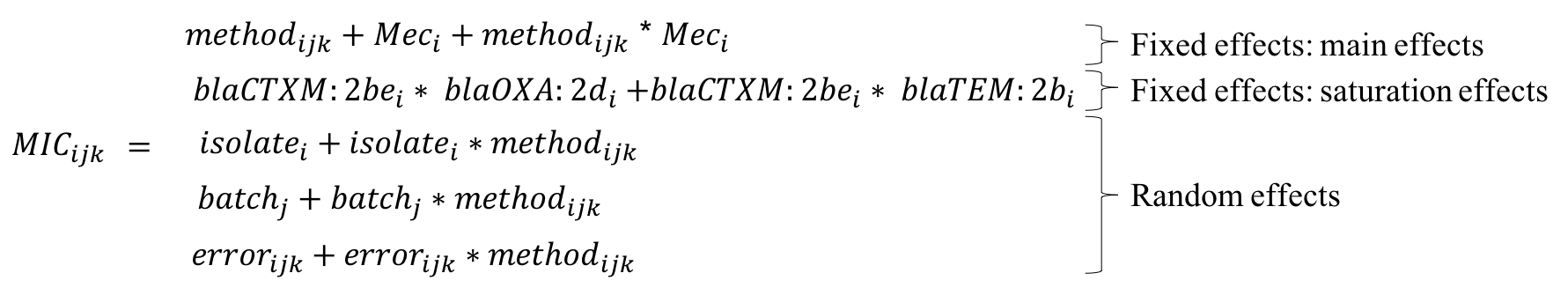

Where for an isolate i, batch j, and test k, and test methods 0 (ratio-based, CLSI), and 1 (fixed-based, EUCAST)

*Mec_i_*

***_=_***

*Other_bla_i +_ blaOXA:2d_i +_ blaCTXM:2be_i +_ blaTEM:2b_i +_ blaSHV:2b_i_*

*_+_ ampCpr_i +_ blaTEMpr_i +_ NFOMP_i_*

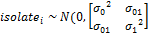

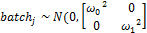

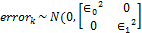

(Note names in equations above represent model components)

*Model-based MIC prediction*

The model above was developed to reflect associations between resistance elements and agar-dilution MIC as accurately as possible, and hence considered potential non-linearity of associations and interactions. However, having developed the model, an obvious question is whether the fixed effect estimates could provide accurate MIC predictions. Cross-validation is a standard method for estimating internal error (i.e. in the same dataset on which models are developed, predicting the observed outcome from the factors included in the model). However, this “training” set was multi-level, including multiple replicates per isolate, potentially with different phenotypes for each of two different testing methodologies (EUCAST and CLSI). To stratify the “training” set for random selection into folds for cross-validation, we therefore first divided the agar dilution subsample isolates into four strata based on each isolate’s upper median MIC for EUCAST and CLSI agar-dilution, namely EUCAST-resistant/CLSI-resistant, EUCAST-resistant/CLSI-intermediate, EUCAST-resistant/CLSI-susceptible and EUCAST- susceptible/CLSI- susceptible. We then split each strata randomly into five folds; fit the final model above on each group of four folds; predicted an MIC for each of EUCAST and CLSI methodologies in the fifth fold; and compared this predicted MIC with the upper median MIC for each isolate for each method. EUCAST MIC predictions agreed with the upper median MIC for 166/261 (64%) isolates and were in essential agreement (within ±1 doubling dilution) for 245/261 (94%). CLSI MIC predictions agreed with the upper median MIC for 155/261 (59%) isolates and were in essential agreement (within ±1 doubling dilution) for 248/261 (95%).

To provide an external estimate of the accuracy of MIC prediction we compared the rounded prediction based on EUCAST fixed estimates with the observed BD Phoenix MIC. As the essential agreement between BD Phoenix and EUCAST agar dilution was 92% we focussed on non-subsample (715/976) isolates for this assessment of external error. Comparisons included all 715 isolates and 704/715 isolates that did not contain either ‘incomplete’ beta-lactamases, or beta-lactamases not present in the agar dilution subsample. Of note, in the 261 subsample isolates EUCAST MIC predictions agreed with BD Phoenix MIC for 175/261 (67%) isolates and were in essential agreement (within ±1 doubling dilution) for 243/261 (93%).

**Supplementary Figures**

**Supplementary Figure S1: Sampling frame and sample selection**

**A: Overall**

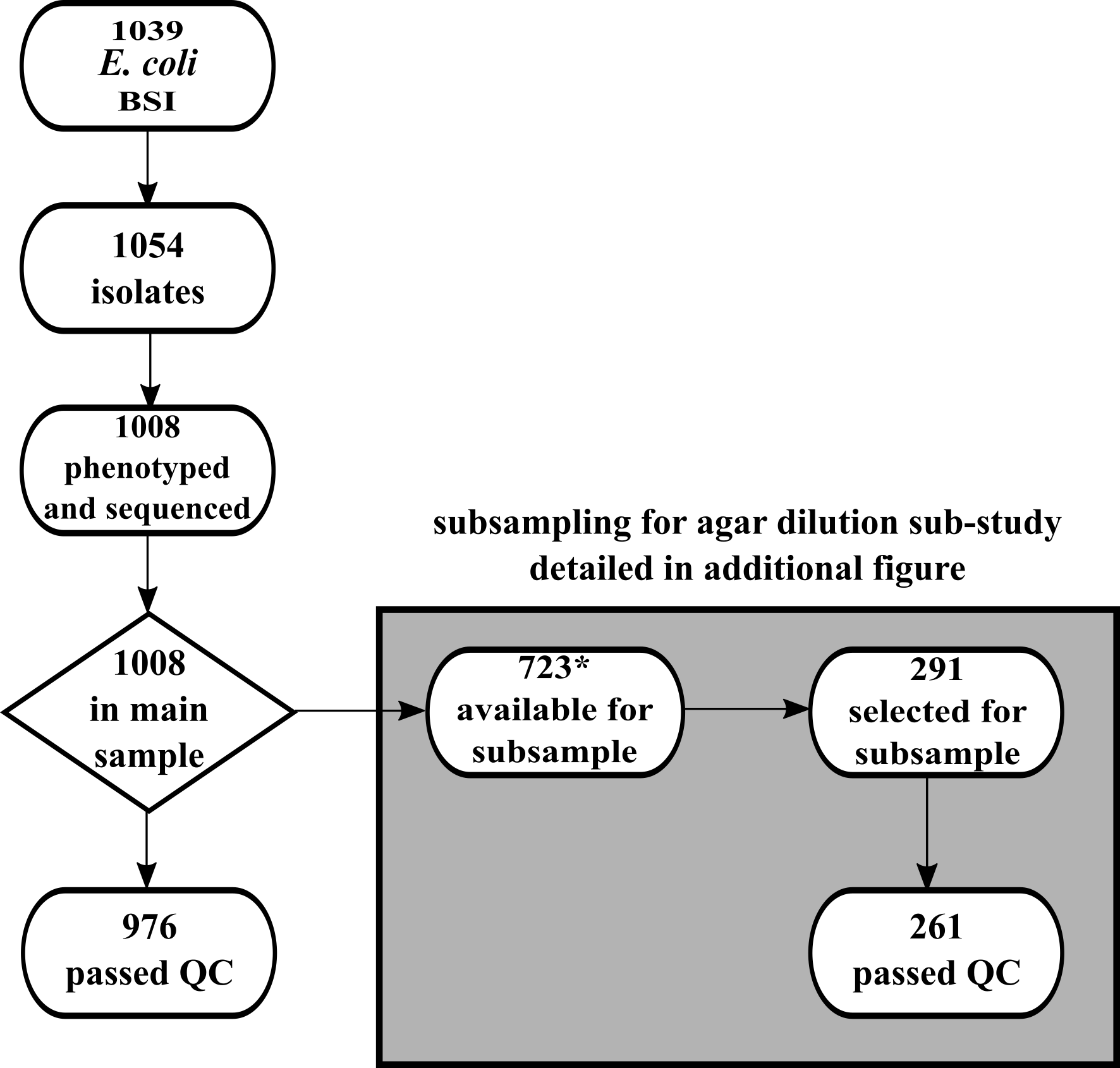

**B: Stratified random sampling strategy for detailed agar dilution phenotyping**

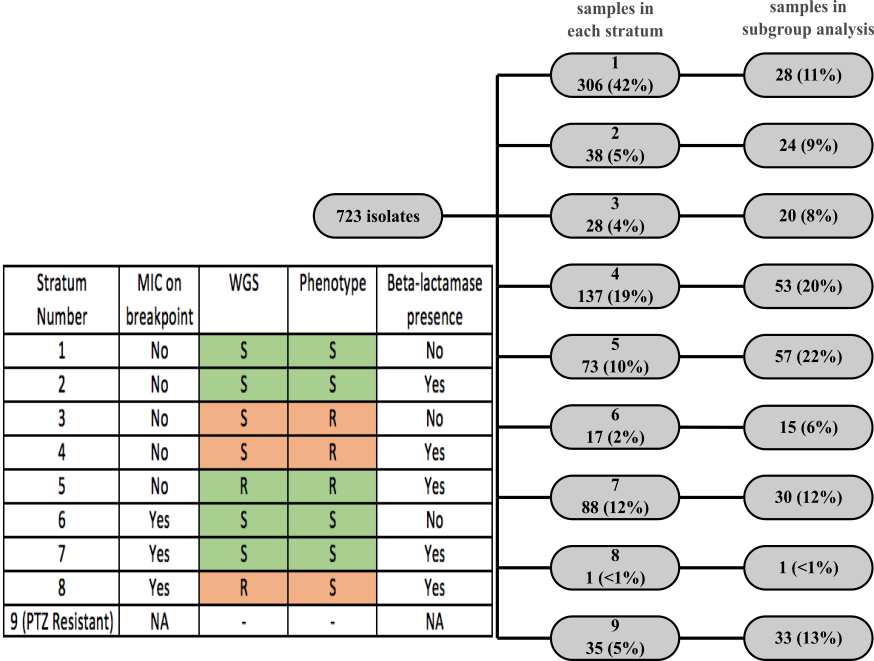

Note: samples included in subsample analysis if in addition to being selected, they passed additional quality control checks (see Supplementary Methods). Orange indicates discordance between WGS-predicted resistance using the basic algorithm (i.e. based on gene presence/absence alone) and AST observed phenotype discordance.

**Supplementary Figure S2: Isolate STs, resistance mechanisms and amoxicillin-clavulanate phenotyping for a) the full dataset N=976) and b) the agar dilution subsample (N=261)**

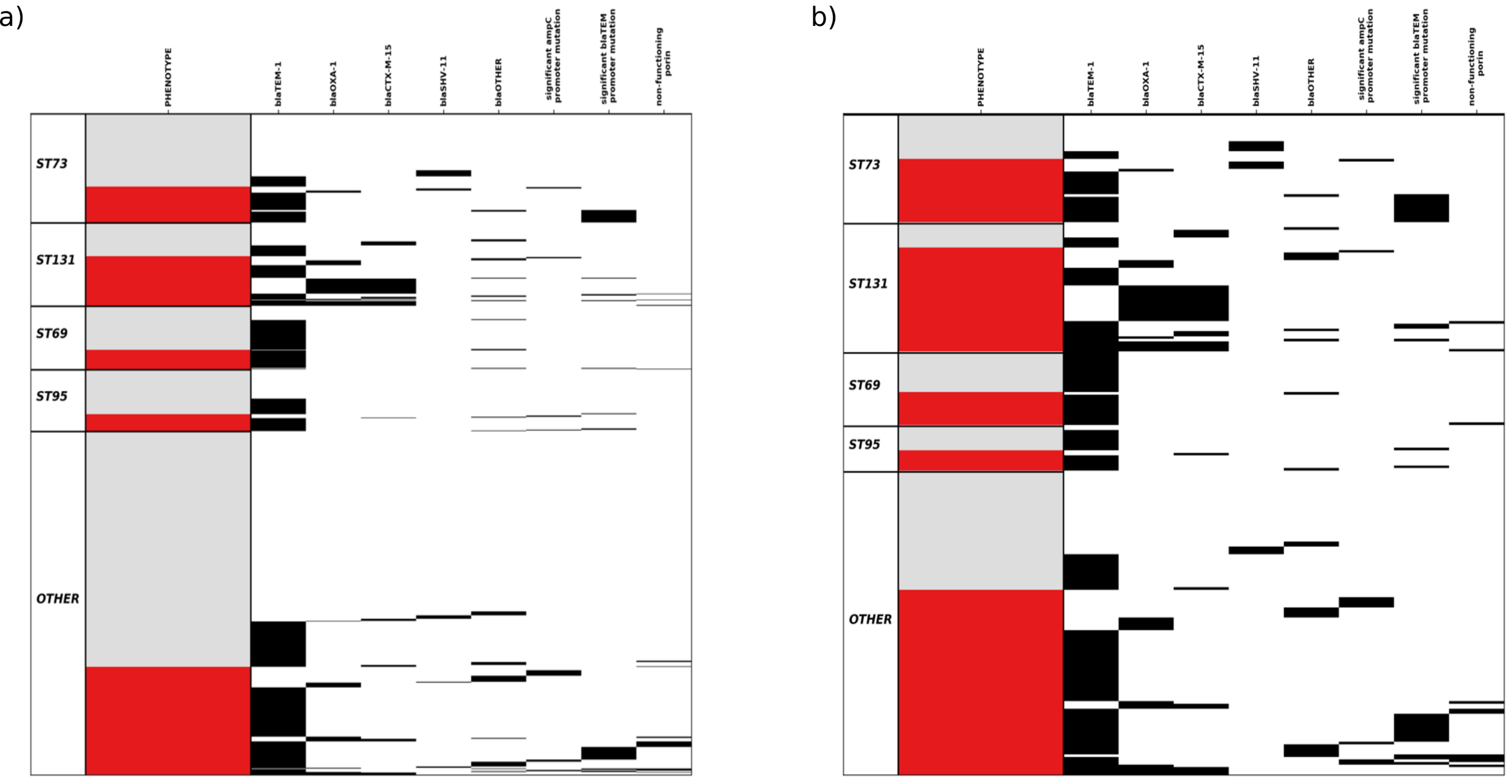

Note: Isolate ST and resistance mechanisms for a) the main dataset (N=976) and b) the agar dilution subsample (N=261). By sampling method, the agar dilution subsample was enriched for amoxicillin-clavulanate resistance. Single horizontal lines represent each isolate. Red indicates resistant by BD Phoenix/EUCAST breakpoints (>8/2 mg/L) and grey susceptible. Black lines indicate the presence of each genetic feature with blaOTHER being any non *bla*_TEM-1_, *bla*_OXA-1_, *bla*_SHV-11_ or *bla*_CTX-M-15_ beta-lactamase (see Supplementary Tables). For promoter mutations/non-functional porin definitions, see Supplementary Methods.

**Supplementary Figure S3: Estimating the coverage copy number cut-off**

**A: ROC curve for selection of DNA copy number threshold for the extended resistance prediction**

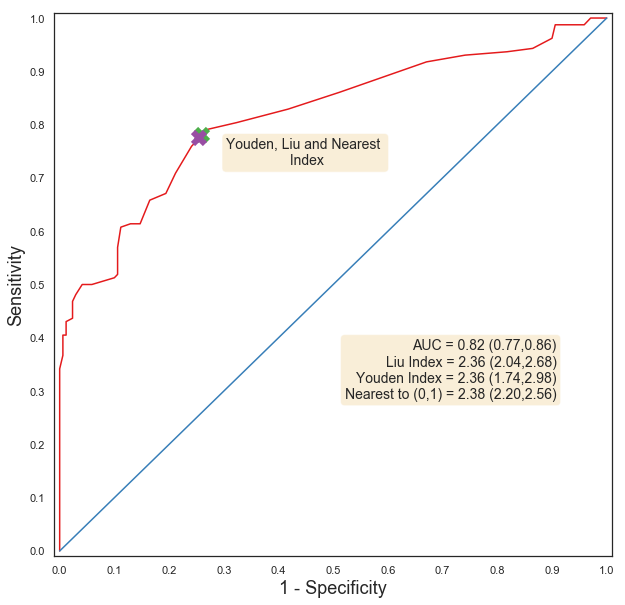

**B: Association between DNA copy number and agar-dilution phenotype in isolates containing only non-inhibitor resistant beta-lactamases**

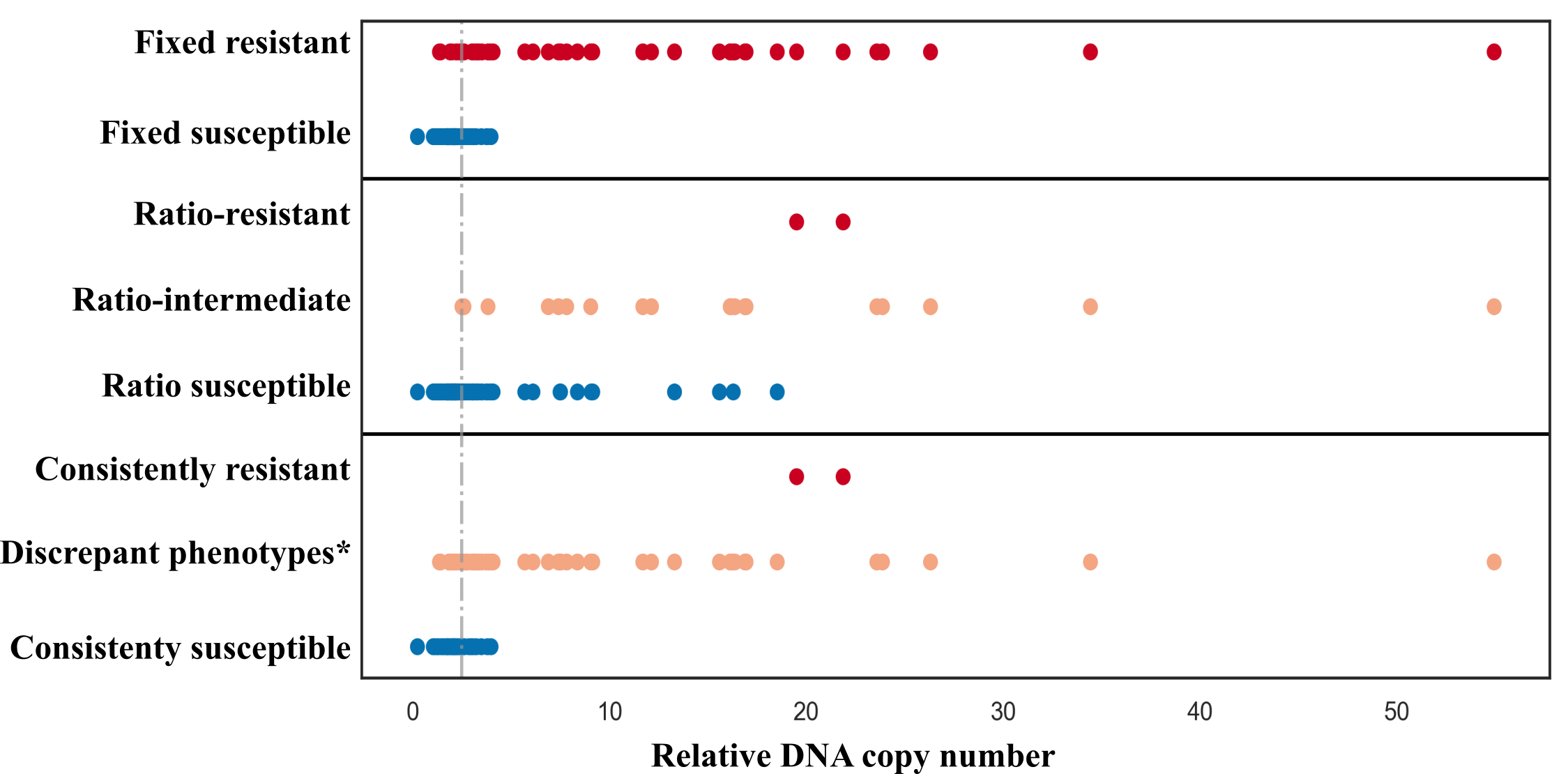

*Non-constant phenotype over repeats, i.e. discrepancy between fixed and ratio phenotypes or discrepancy with fixed/ratio repeats.

Note: including the relative copy number of beta-lactamases in 107 isolates from the agar dilution subsample which each contained only one non-inhibitor resistant beta-lactamase (excluding one isolate which contained a novel *bla*_CTX-M_-like gene with a SDN mutation, see main text results/discrepancies). Genes included 99 non-inhibitor resistant *bla*_TEM_ genes, 10 non-inhibitor resistant *bla*_SHV_ genes and 6 *bla*_CTX-M_ genes. The grey dotted line represents the predefined coverage cutoff threshold of 2.5x. Colors highlight different phenotypes as denoted on the left of the panel.

**C: MICs in isolates containing only low copy number bla_TEM-1_**

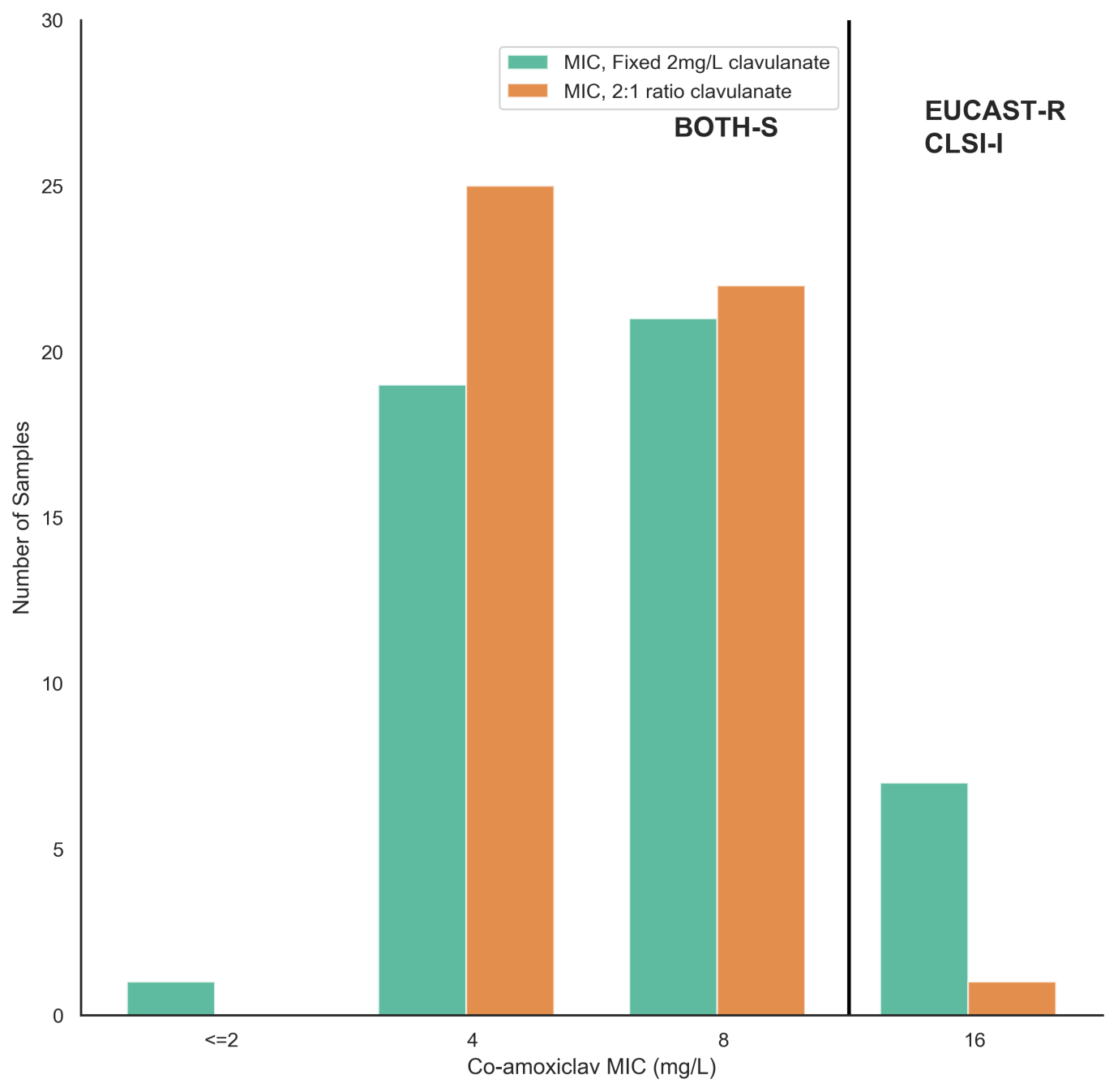

### **Supplementary Figure S4: Maximum MIC doubling dilution difference across repeats by method for subsample isolates**

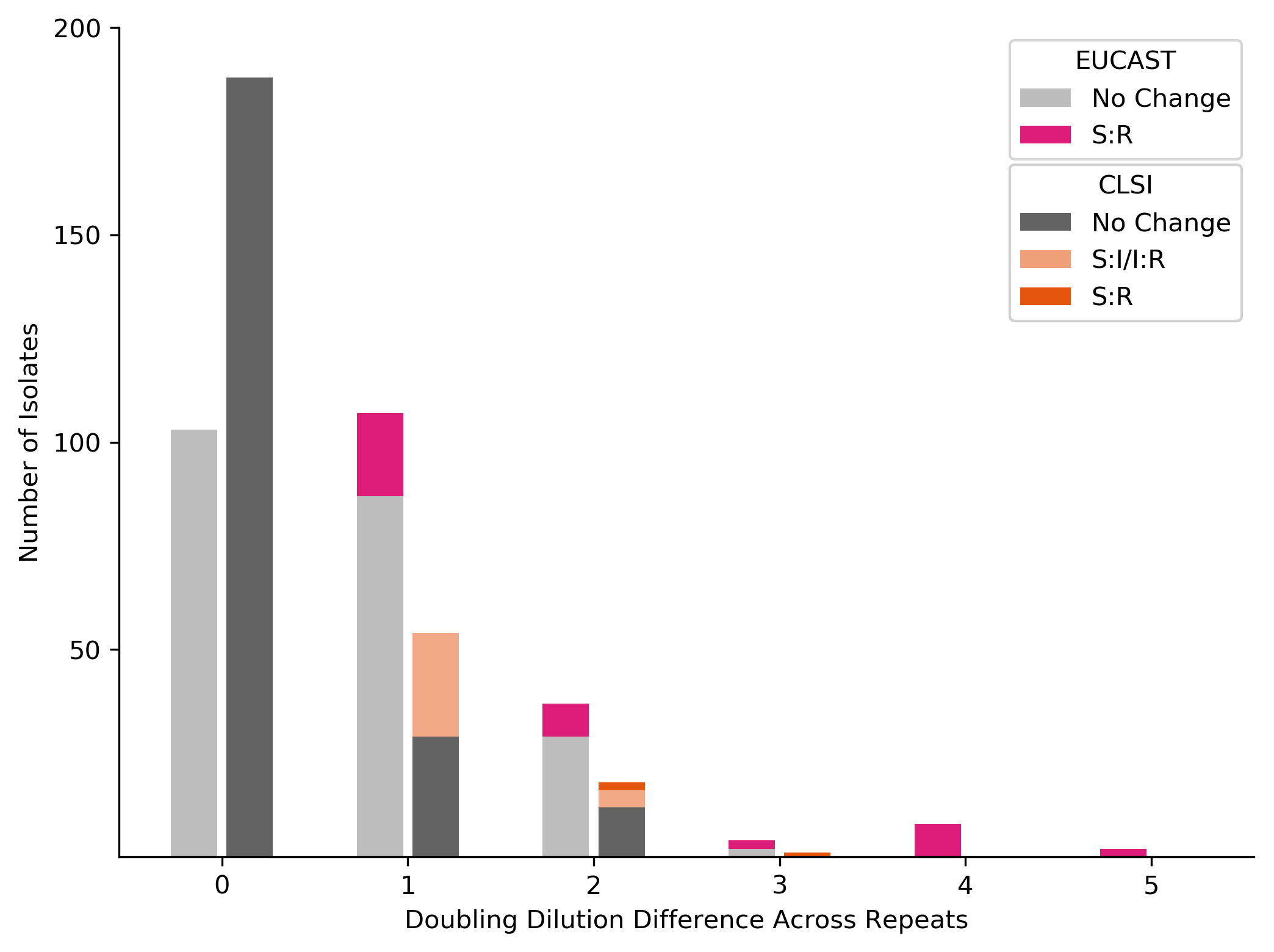

**Supplementary Tables**

**Supplementary Table S1: Beta-lactam antibiograms from automated susceptibility testing in the Oxford University Hospitals microbiology laboratory**

|  | **Number (%) of resistant isolates** | | | |
| --- | --- | --- | --- | --- |
|  | **Overall** | | **Agar dilution subsample** | |
| **Antibiotic** | **All isolates (N=976)** | **Amoxicillin-clavulanate-resistant (N=339)** | **All isolates**  **(N=261)** | **Amoxicillin-clavulanate-resistant (N=160)** |
| **Amoxicillin-clavulanate** | 339 (35%) | 339 (100%) | 160 (61%) | 160 (100%) |
| **Ampicillin** | 531 (54%) | 332 (98%) | 219 (84%) | 159 (99%) |
| **Aztreonam** | 61 (6%) | 44 (13%) | 31 (12%) | 24 (15%) |
| **Cefepime­­** | 57 (7%) | 44 (13%) | 32 (12%) | 25 (16%) |
| **Cefoxitin *** | 30 (3%) | 28(8%) | 19 (7%) | 19 (12%) |
| **Ceftazidime** | 53 (5%) | 42 (12%) | 32 (12%) | 27 (17%) |
| **Ceftriaxone** | 79 (8%) | 53 (16%) | 38 (15%) | 31 (19%) |
| **Ertapenem** | 3 (0.3%) | 2 (0.2%) | 1 (0.3%) | 1 (0.6%) |
| **Meropenem** | 0 (0%) | 0 (0%) | 0 (0%) | 0 (0%) |
| **Piperacillin-tazobactam** | 50 (5%) | 50 (15%) | 34 (13%) | 34 (21%) |

* Number of isolates positive for ampC production screen

**Supplementary Table S2: Beta-lactamases**

**A: Identified beta-lactamases and amoxicillin-clavulanate phenotype**

| **Feature** | | **Mutations**  **(Ambler positions)** | **Number of isolates with mechanism (total sequenced [n=976])** | **Number of isolates with only this mechanism (total sequenced [n=976])** | **Number of isolates amoxicillin-clavulanate resistant by BD Phoenix** | |
| --- | --- | --- | --- | --- | --- | --- |
|  |  |  |  |  | **Total (% of no with this mechanism)** | **Not beta-lactam resistant by any other^o^ genetic mechanism** |
| ***bla*_TEM_ beta-lactamases** | | | | | | |
|  | TEM-1 |  | 409 | 312 | 244 (60%) | 158 (50%) |
|  | TEM-1-like* | Ser243Ala | 1 | 1 | 1 (100%) | 1 (100%) |
|  | TEM-30 |  | 4 | 3 | 4(100%) | 3 (100%) |
|  | TEM-31 |  | 2 | 0 | 1 (100%) | - |
|  | TEM-33 |  | 1 | 1 | 2(100%) | 1 (100%) |
|  | TEM-40 |  | 5 | 2 | 5(100%) | 2 (100%) |
|  | TEM-135 |  | 1 | 1 | 1 (100%) | 1 (100%) |
|  | TEM-190 |  | 1 | 1 | 1 (100%) | 1 (100%) |
|  | TEM-unknown* |  | 3 | 3 | 2 (67%) | 2 (67%) |
| ***bla*_CTX-M-1_ group beta-lactamases** | | | | | | |
|  | CTX-M-1 |  | 3 | 3 | 0 (0%) | 0 (0%) |
|  | CTX-M-15 |  | 52 | 9 | 40 (77%) | 0 (0%) |
|  | CTX-M-15-like | Ser130Gly | 1 | 1 | 1 (100%) | 1 (100%) |
|  | CTX-M-3 |  | 1 | 0 | 1 (100%) | -^+^ |
|  | CTX-M-55 |  | 2 | 0 | 1 (100%) | - |
|  | CTX-M-15-like | Ser130Gly | 1 | 1 | 1 (100%) | 1 (100%) |
| ***bla*_CTX-M-9_ group beta-lactamases** | | | | | | |
|  | CTX-M-9 |  | 1 | 0 | 1 (100%) | -^+^ |
|  | CTX-M-14b |  | 9 | 1 | 5 (56%) | 1 (20%) |
|  | CTX-M-27 |  | 4 | 3 | 2 (50%) | 0 (0%) |
| **Other beta-lactamases identified** | | | | | | |
|  | OXA-1 |  | 60 | 17 | 60 (100%) | 17 (100%) |
|  | OXA-48 |  | 1 | 0 | 1 (100%) | - |
|  | SHV-11 |  | 20 | 18 | 6 (30%) | 4 (22%) |
|  | SHV-12 |  | 3 | 1 | 1 (33%) | 0 (0%) |
|  | CMY-2 |  | 5 | 1 | 5 (100%) | 1 (100%) |
|  | CMY-60 |  | 1 | 1 | 0 (0%) | 0 (0%) |
|  | OXA-unknown |  | 1 | 1 | 0 (0%) | 0 (0%) |
|  | CMY-unknown |  | 2 | 1 | 1 (50%) | 0 (0%) |
|  | LAT-unknown |  | 1 | 0 | 1 (100%) | - |

o: Other beta-lactam resistance mechanisms includes any other beta-lactamase, ampC promoter mutations suggestive of ampC hyper production and functional porin loss

-: no isolates in this category

*: “like” indicates inexact match with parent reference sequence, “unknown” indicates inability to identify a complete gene (ARIBA flag),

**B: Frequencies and combinations of multiple beta-lactamases seen in isolates**

| Beta-lactamase combination | Number of isolates with combination |
| --- | --- |
| *bla*_CTX-M-15_, *bla*_OXA-1_ | 27 |
| *bla*_CTX-M-15_, *bla*_OXA-1_, *bla*_TEM-1_ | 10 |
| *bla*_CTX-M-14_, *bla*_TEM-1_ | 8 |
| *bla*_CTX-M-15_, *bla*_TEM-1_ | 6 |
| *bla*_OXA-1_, *bla*_TEM-1_ | 3 |
| *bla*_CMY-2_, *bla*_TEM-1_ | 2 |
| *bla*_SHV-11_, *bla*_TEM-1_ | 2 |
| *bla*_SHV-12_, *bla*_TEM-1_ | 2 |
| *bla*_CMY-2_, *bla*_LAT*_, *bla*_TEM-1_ | 1 |
| *bla*_CMY-2_, *bla*_CMY*_, *bla*_TEM-1_ | 1 |
| *bla*_CTX-M-3_, *bla*_TEM-40_ | 1 |
| *bla*_CTX-M-9_, *bla*_TEM-1_ | 1 |
| *bla*_CTX-M-27_, *bla*_TEM-1_ | 1 |
| *bla*_CTX-M-55_, *bla*_OXA-1_ | 1 |
| *bla*_OXA-48_, *bla*_TEM-1_ | 1 |

**Supplementary Table S3:** **Promoter sequences**

**A: *ampC* promotors**

| **Mutation pattern** | **Number of isolates with mutation pattern (total sequenced [n=976])** | **Number of isolates with mutation pattern and no beta-lactamases** | **Number of isolates amoxicillin-clavulanate resistant by BD Phoenix (% of total with this mechanism)** | |
| --- | --- | --- | --- | --- |
|  |  |  | **Total** | **Not beta-lactam resistant by any other genetic mechanism^o^** |
| Nil | 368 | 156 | 120 (33%) | 0 (0%) |
| -28 G>A | 168 | 55 | 83 (49%) | 0 (0%) |
| -28 G>A\|17 C>T | 168 | 93 | 39 (23%) | 1 (1%) |
| 58 C>T | 125 | 74 | 36 (29%) | 0 (0%) |
| -18 G>A\|-1 C>T\|58 C>T | 69 | 32 | 21 (45%) | 1 (3%) |
| 22 C>T\|26 T>G\|27 A>T\|32 G>A | 17 | 7 | 9 (53%) | 1 (14%) |
| -28 G>A \|58 C>T | 9 | 3 | 1 (11%) | 0 (0%) |
| 48 A>G\|58 C>T | 4 | 4 | 0 (0%) | 0 (0%) |
| **-42 C>T***\|-18 G>A\|-1 C>T\|58 C>T | 3 | 1 | 3 (100%) | 1 (100%) |
| -18 G>A\|-11 C>T\|-1 C>T\|58 C>T | 3 | 1 | 3 (100%) | 1 (100%) |
| 37 G>T | 3 | 2 | 1 (33%) | 0 (0%) |
| **-32 T>A** | 3 | 3 | 3 (100%) | 3 (100%) |
| **-32 T>A**\|58 C>T | 3 | 0 | 3 (100%) | - |
| -28 G>A\|-**24_-23InsT** \|17 C>T | 2 | 2 | 2 (100%) | 2 (100%) |
| 34_35InsC\|58 C>T | 2 | 2 | 0 (0%) | 0 (0%) |
| -28 G>A \|**-15_-14InsT**\|17 C>T | 2 | 2 | 2 (100%) | 2 (100%) |
| -28 G>A \|-10_1Del\|17 C>T | 2 | 2 | 0 (0%) | 0 (0%) |
| 33 G>A | 2 | 2 | 0 (0%) | 0 (0%) |
| **-15_-14InsG** | 1 | 1 | 1 (100%) | 1 (100%) |
| -12_-5Del\|58 C>T | 1 | 1 | 1 (100%) | 1 (100%) |
| -12_-5Del | 1 | 1 | 0 (0%) | 0 (0%) |
| -9_-8Del\|58 C>T | 1 | 1 | 0 (0%) | 0 (0%) |
| -11 C>T | 1 | 1 | 1 (100%) | 1 (100%) |
| -29 C>T | 1 | 1 | 0 (0%) | 0 (0%) |
| **-42 C>T\|**-18 G>A\|-1 C>T\|20_21InsG\|58 C>T | 1 | 1 | 1 (100%) | 1 (100%) |
| 20DelC | 1 | 1 | 0 (0%) | 0 (0%) |
| -28 G>A \|**-15_-14InsGT**\|17 C>T \|41 T>A | 1 | 1 | 1 (100%) | 1 (100%) |
| -28 G>A \|-12_-5Del\|17 C>T | 1 | 1 | 0 (0%) | 0 (0%) |
| -28 G>A \|**-16_-15InsC** | 1 | 1 | 1 (100%) | 1 (100%) |
| -28 G>A \|-9_2Del\|17 C>T | 1 | 1 | 0 (0%) | 0 (0%) |
| -28 G>A \|17 C>T \|20 C>T | 1 | 1 | 0 (0%) | 0 (0%) |
| -28 G>A \|33 G>T | 1 | 1 | 1 (100%) | 1 (100%) |
| -28 G>A \|34 G>A\|58 C>T | 1 | 0 | 1 (100%) | - |
| -28 G>A \|26-28Del | 1 | 1 | 1 (100%) | 1 (100%) |
| 30 G>A | 1 | 0 | 0 (0%) | 0 (0%) |
| **-32 T>A**\|-28 G>A | 1 | 1 | 1 (100%) | 1 (100%) |
| **-32 T>A**\|-28 G>A \|**-21_-20InsT**\|-11 C>T\|17 C>T | 1 | 1 | 1 (100%) | 1 (100%) |
| **-32 T>A**\|-28 G>A \|17 C>T | 1 | 0 | 1 (100%) | - |
| Promoter not identified/unable to assemble | 3 | 3 | 2 (66%) | 2 (66%) |

o: Other beta-lactam resistance mechanisms includes any other beta-lactamase, *ampC* promoter mutations suggestive of ampC hyper production and functional porin loss

-: no isolates in this category , *:letters in bold represent pre-defined significant mutations.

**B: amoxicillin-clavulanate MICs of *ampC* hyper-producing isolates**

| Isolate | Amoxicillin-clavulanate MIC | | Other beta-lactam resistance mechanisms |
| --- | --- | --- | --- |
|  | Median fixed MIC (mg/L) | Median ratio MIC (mg/L) |  |
| ampC_1 | >128/2 | 32/16 | Low copy number *bla*_TEM-1_ |
| ampC_2 | 128/2 | 32/16 |  |
| ampC_3 | 128/2 | 32/16 |  |
| ampC_4 | >128/2 | 64/32 | Low copy number *bla*_TEM-1_ |
| ampC_5 | 64/2 | 32/16 |  |
| ampC_6 | 128/2 | 32/16 | Low copy number *bla*_TEM-1_ |
| ampC_7 | 128/2 | 32/16 |  |
| ampC_8 | 128/2 | 32/16 |  |
| ampC_9 | 128/2 | 32/16 |  |

**C: *bla*_TEM_ promotors**

| **Mutation pattern** | **Number of isolates with mutation pattern (total isolates with blaTEM [n=427])** | **Closest promoter type^x^** | **Number of isolates amoxicillin-clavulanate resistant by BD Phoenix (% of total with this mechanism)** |
| --- | --- | --- | --- |
| Nil | 254 | P3 | 122 (48%) |
| 175 G>A | 96 | P3 | 68 (71%) |
| **32 C>T*** | 26 | Pa/Pb | 26 (100%) |
| **32 C>T**\|175 G>A | 15 | Pa/Pb | 15 (100%) |
| **162 G>T** | 4 | P4 | 4 (100%) |
| **162 G>T**\|175 G>A | 2 | P4 | 1 (50%) |
| 43 G>A\|175 G>A | 1 | P3 | 0 (0%) |
| **30_38Del** | 1 | Pc/Pd | 1 (100%) |
| **30_38Del**\|175 G>A | 1 | Pc/Pd | 1 (100%) |
| 65 G>A\|175 G>A | 1 | P3 | 0 (0%) |
| Promoter not identified/unable to assemble | 26 | NA | 13 (50%) |

*: Letters in bold represent mutations we had pre-defined as significant.

x: For promoter type and the effects of mutations, see Lartigue et al. Antimicrobial agents and chemotherapy. 2002;46(12):4035-7

**Supplementary Table S4: Disrupted porin genes**

**A: Beta-lactam resistance context and antibiogram for isolates with disrupted ompF porins**

| **Isolate** | **Extended algorithm prediction** | **Porin gene issue^o^** | **Other beta-lactam mechanisms** | **Minimum inhibitory concentration (mg/L)** | | | | | | |
| --- | --- | --- | --- | --- | --- | --- | --- | --- | --- | --- |
|  |  |  |  | **Ampicillin** | **Co-amoxiclav** | **Piperacillin-tazobactam** | **Cefoxitin** | **Ceftriaxone** | **Ceftazidime** | **Meropenem** |
| **AMC_43** | R | truncation | De-repressed chromosomal *ampC*, *bla*_TEM-1_ | >8 | >32/2 | 8/4 | >16 | 1 | 8 | <=0.5 |
| **AMC_690** | R | truncation | De-repressed chromosomal *ampC* | >8 | >32/2 | <=4/4 | >16 | 1 | 4 | <=0.5 |
| **AMC_113** | R | truncation | *bla*_CTX-M-15_, *bla*_OXA-1_ | >8 | >32/2 | 32/4 | <=4 | >4 | >16 | <=0.5 |
| **AMC_884** | R | truncation | *bla*_OXA-1,_ *bla*_TEM-1_ | >8 | >32/2 | >64/4 | <=4 | <=1 | <=1 | <=0.5 |
| **AMC_163** | R | truncation | *bla*_OXA-1_ | >8 | >32/2 | 16 | 16 | <=1 | <=1 | <=0.5 |
| **AMC_778** | R | truncation | *bla*_TEM-30_ | >8 | >32/2 | <=4/4 | 8 | <=1 | <=1 | <=0.5 |
| **AMC_173** | R | truncation | High copy number *bla*_TEM-1_ | >8 | 8/2 | <=4/4 | <=4 | <=1 | <=1 | <=0.5 |
| **AMC_211** | R | truncation | High copy number *bla*_TEM-1_ | >8 | 32/2 | <=4/4 | 16 | <=1 | <=1 | <=0.5 |
| **AMC_366** | R | truncation | High copy number *bla*_TEM-1_ | >8 | 8/2 | <=4/4 | <=4 | <=1 | <=1 | <=0.5 |
| **AMC_628** | S | truncation | - | <=2 | 4/2 | <=4/4 | <=4 | <=1 | <=1 | <=0.5 |
| **AMC_280** | S | truncation | - | <=2 | 4/2 | <=4/4 | <=4 | <=1 | <=1 | <=0.5 |
| **AMC_131** | S | truncation | - | 4 | 8/2 | <=4/4 | >16 | <=1 | <=1 | <=0.5 |
| **AMC_651** | R | assembly issue | *bla*_CTX-M-15_, high copy number *bla*_TEM-1_ | >8 | 8/2 | <=4/4 | <=4 | >4 | 8 | <=0.5 |
| **AMC_63** | R | assembly issue | High copy number *bla*_TEM-1_ | >8 | >32/2 | <=4/4 | 8 | <=1 | <=1 | <=0.5 |
| **AMC_311** | R | assembly issue | High copy number *bla*_TEM-1_ | >8 | 16/2 | <=4/4 | <=4 | <=1 | <=1 | <=0.5 |
| **AMC_258** | R | assembly issue | High copy number *bla*_TEM-1_ | >8 | 32/2 | <=4/4 | 8 | <=1 | <=1 | <=0.5 |

*: Isolates predicted co-amoxiclav resistant due to non-functional porin in the presence of (any) beta-lactamase

o: Truncations occur when frameshift mutations leading to premature stop codons are found in the sequence. Assembly issues are when our search method (ARIBA) failed to completely and uniquely assemble the gene (see Methods) or the gene was only found at <10x coverage (which was typically associated with multiple frameshift mutations, suggesting uncertainty of the sequence).

### Note: Genetic data for each isolate are available as part of PRJNA540750 at NCBI. Isolate identifiers are as above.

### **B: Beta-lactam resistance context and antibiogram for isolates with disrupted ompC porins**

| **Isolate** | **Extended algorithm prediction** | **Porin gene issue^o^** | **Other beta-lactam mechanisms** | **Minimum inhibitory concentration (mg/L)** | | | | | | |
| --- | --- | --- | --- | --- | --- | --- | --- | --- | --- | --- |
|  |  |  |  | **Ampicillin** | **Amoxicillin-clavulanate** | **Piperacillin-tazobactam** | **Cefoxitin** | **Ceftriaxone** | **Ceftazidime** | **Meropenem** |
| **AMC_116** | R | truncation | *bla*_CTX-M-15_, *bla*_OXA-1_, high copy number *bla*_TEM-1_ | >8 | 32/2 | <=4/4 | 8 | >4 | 1 | <=0.5 |
| **AMC_229** | R | truncation | High copy number *bla*_TEM-1_ | >8 | >32/2 | <=4/4 | <=4 | <=1 | <=1 | <=0.5 |
| **AMC_824*** | R | truncation | *bla*_TEM-1_ | >8 | >32/2 | <=4/4 | <=4 | <=1 | <=1 |  |
| **AMC_264** | S | truncation | - | 4 | 8/2 | <=4/4 | 8 | <=1 | <=1 | <=0.5 |
| **AMC_188** | S | truncation | - | <=2 | 4/2 | <=4/4 | <=4 | <=1 | <=1 | <=0.5 |
| **AMC_562** | S | truncation | - | <=2 | <=2/2 | <=4/4 | <=4 | <=1 | <=1 | <=0.5 |
| **AMC_799** | R | assembly issue | *bla*_OXA-1_ | >8 | >32/2 | 64/4 | 8 | <=1 | 1 | <=0.5 |
| **AMC_805** | R | assembly issue | High copy number *bla*_TEM-1_ with a non-P3 *bla*_TEM_ promoter | >8 | >32/2 | >64/4 | >16 | 1 | 2 |  |
| **AMC_86** | R | assembly issue | High copy number *bla*_TEM-1_ | >8 | >32/2 | <=4/4 | <=4 | <=0.5 | <=0.5 | <=0.5 |
| **AMC_121** | R | assembly issue | High copy number *bla*_TEM-1_ | >8 | >32/2 | <=4/4 | <=4 | <=1 | <=1 | <=0.5 |
| **AMC_228** | R | assembly issue | High copy number *bla*_TEM-1_ | >8 | 16/2 | <=4/4 | <=4 | <=1 | <=1 | <=0.5 |
| **AMC_412*** | R | assembly issue | *bla*_TEM-1_ | >8 | 16/2 | <=4/4 | <=4 | <=1 | <=1 | <=0.5 |
| **AMC_596** | S | assembly issue | - | <=2 | <=2/2 | <=4/4 | <=4 | <=1 | <=1 | <=0.5 |
| **AMC_696** | S | assembly issue | - | <=2 | <=2/2 | <=4/4 | <=4 | <=1 | <=1 | <=0.5 |
| **AMC_929** | S | assembly issue | - | <=2 | <=2/2 | <=4/4 | <=4 | <=1 | <=1 | <=0.5 |

*: Isolates predicted co-amoxiclav resistant due to non-functional porin in the presence of (any) beta-lactamase

o: Truncations occur when frameshift mutations leading to premature stop codons are found in the sequence. Assembly issues are when our search method (ARIBA) failed to completely and uniquely assemble the gene (see Methods) or the gene was only found at <10x coverage (which was typically associated with multiple frameshift mutations, suggesting uncertainty of the sequence).

Note: Genetic data for each isolate are available as part of PRJNA540750 at NCBI. Isolate identifiers are as above.

**Supplementary Table S5: Beta-lactamase classification for random-effects models for log2MIC, agar dilution subsample only**

| **Model Category** | **Number of isolates**  **with this category (% of total 261)** | **Included components** |
| --- | --- | --- |
| **blaTEM:2b** | 155 (59%) | *bla*_TEM-1_ (N=154), *bla*_TEM-135_ (N=1) |
| **blaCTXM:2be** | 37 (14%) | *bla*_CTX-M-15_ (N=30), *bla*_CTX-M-14b_ (N=2), *bla*_CTX-M-1_ (N=2), *bla*_CTX-M-27_ (N=2), *bla*_CTX-M-3_ (N=1) |
| **blaOXA:2d** | 35 (13%) | *bla*_OXA-1_ (N=35) |
| **blaSHV:2b** | 10 (4%) | *bla*_SHV-11_ (N=10) |
| **Other_bla*** | 13 (5%) | *bla*_TEM-40_ (N=2), *bla*_TEM-30_(N=3), *bla*_CMY-2_ (N=3), *bla*_OXA-48_ (N=1), *bla*_TEM-190_ (N=1)*, bla*_TEM-33_ (N=1)  3 non-complete beta-lactamases |

* : one isolate contained both blaCMY-2 and a non-complete beta-lactamase

Note: see Supplementary Table S2A for individual beta-lactamases identified across all study isolates

**Supplementary Table S6: Unadjusted effect of resistance features on agar dilution log2 MIC from univariable random-effects models**

| **Element** | **Testing method** | **Mean log_2_(MIC)**  **(95% CI)** | **log_2_(MIC) change**^†^ | **Corresponding MIC range covering the 95% CI (mg/L)*** |
| --- | --- | --- | --- | --- |
| **No element/feature** | CLSI | 2.08 (1.77, 2.40) | - | 2-8 |
|  | EUCAST | 2.10 (1.70, 2.51) | - | 2-8 |
| **blaTEM:2b** | CLSI | 3.32 (3.17, 3.48) | 1.25 (0.93,1..56) | 8-16 |
|  | EUCAST | 4.30 (4.03, 4.57) | 2.20 (1.67, 2.73) | 16-32 |
| **blaOXA:2d** | CLSI | 3.74 (3.57, 3.92) | 1.66 (1.27,2.52) | 8-16 |
|  | EUCAST | 5.20 (4.91, 5.49) | 3.10 (2.57,3.62) | 16-64 |
| **blaCTXM:2be** | CLSI | 3.49 (3.24, 3.73) | 1.40 (0.99,1.82) | 8-16 |
|  | EUCAST | 4.68 (4.20, 5.15) | 2.97 (1.96,3.18) | 16-64 |
| **blaSHV:2b** | CLSI | 2.30 (1.95, 2.65) | 0.22 (-0.49,0.92) | 2-8 |
|  | EUCAST | 2.50 (1.73, 3.27) | 0.40 (-0.54,1.33) | 2-16 |
| **Other_bla^o^** | CLSI | 4.53 (4.01,5.07) | 2.46 (1.80,3.11) | 16-64 |
|  | EUCAST | 6.69 (5.75,7.62) | 4.59 (3.69, 5.48) | 128-256 |
| **ampC promoter mutation** | CLSI | 5.11 (4.85,5.37) | 3.02 (2.29,3.77) | 16-64 |
|  | EUCAST | 7.11 (6.65,7.57) | 5.01 (4.05,5.96) | 128-256 |
| **blaTEM promoter mutation** | CLSI | 4.27 (4.01,4.52) | 2.18 (1.74,2.63) | 16-32 |
|  | EUCAST | 6.00 (5.58,6.42) | 3.90 (3.30,4.49) | 64-128 |
| **Suspected loss of function in ompC/ompF** | CLSI | 4.36 (3.61,5.11) | 2.28 (2.55,3.01) | 8-64 |
|  | EUCAST | 6.27 (5.27,7.27) | 4.17 (3.22,5.11) | 64-256 |

^o^: Other beta-lactamases, *bla*_TEM-40_ (N=2), *bla*_TEM-30_(N=3), *bla*_CMY-2_ (N=3), *bla*_OXA-48_ (N=1), *bla*_TEM-190_ (N=1)*, bla*_TEM-33_ (N=1),3 non-complete beta-lactamases

*: MIC interpretation of the 95% CI

^†^: log_2_(MIC) change relative to the no-element/feature for each test method. E.g. CLSI log_2_(MIC) change for CLSI is expressed relative to the CLSI no-element/feature MIC

**Supplementary Table S7: Results from multivariable random-effects model**

### **A: Independent effects of resistance features on agar dilution log2 MIC from random-effects models**

| **Element Type** | **Element*** | **Modelled as** | **Test method** | **Change in log_2_(MIC) (95% CI)** | **P** | **Heterogeneity between CLSI and EUCAST** |
| --- | --- | --- | --- | --- | --- | --- |
| **Overall** |  |  |  |  |  |  |
|  | - |  | CLSI | - | - |  |
|  |  |  | EUCAST | 0.09 (-0.08,0.26) | 0.29 |  |
| **Transmissible** |  |  |  |  |  |  |
|  | blaTEM:2b | Presence/absence | CLSI | 0.36 (0.08,0.65) | 0.01 | 0.05 |
|  |  |  | EUCAST | 0.14  (-0.27,0.54) | 0.51 |  |
|  | blaTEM:2b | Per doubling of copy number | CLSI | 0.38 (0.29,0.48) | <0.0001 | <0.0001 |
|  |  |  | EUCAST | 0.79 (0.66,0.93) | <0.0001 |  |
|  | blaCTXM:2be | Presence/absence | CLSI | 0.88 (0.42,1.35) | <0.0001 | 0.47 |
|  |  |  | EUCAST | 0.97 (0.40,1.55) | 0.001 |  |
|  | blaOXA:2d | Presence/absence | CLSI | 1.61 (1.17,2.05) | <0.0001 | <0.0001 |
|  |  |  | EUCAST | 2.74 (2.17,3.30) | <0.0001 |  |
|  | blaSHV:2b | Presence/absence | CLSI | 0.03  (-0.71,0.78) | 0.93 | 0.03 |
|  |  |  | EUCAST | -0.61  (-1.68,0.46) | 0.27 |  |
|  | blaSHV:2b* | Per doubling of copy number | CLSI | 0.28  (-0.05,0.60) | 0.10 | 0.001 |
|  |  |  | EUCAST | 0.71 (0.23,1.18) | 0.004 |  |
|  | Other_bla | Presence/absence | CLSI | 2.12 (1.70,2.54) | <0.0001 | <0.0001 |
|  |  |  | EUCAST | 3.79 (3.17,4.40) | <0.0001 |  |
| **Non-transmissible** | |  |  |  |  |  |
|  | ampC promoter mutation (ampCpr) | Presence/absence | CLSI | 2.60 (2.09,3.11) | <0.0001 | <0.0001 |
|  |  |  | EUCAST | 4.25 (3.52,4.99) | <0.0001 |  |
|  | Suspected loss of function in ompC/ompF (NFOMP) | Presence/absence | CLSI | 0.26  -0.21,0.72) | 0.27 | 0.04 |
|  |  |  | EUCAST | 0.63  (-0.03,1.30) | 0.06 |  |
| **Additional Interactions** |  |  |  |  |  |  |
|  | TEM promoter mutation | Presence/absence | CLSI | 1.27 (0.97,1.56) | <0.0001 | <0.0001 |
|  | (blaTEMpr) |  | EUCAST | 2.43 (2.00,2.86) | <0.0001 |  |
|  | CTX-MxTEM | Both present |  | -0.88  (-1.40,-0.35) | 0.001 | - |
|  | CTX-MxOXA | Both present |  | -0.62 (-1.26,0.01) | 0.05 | - |

* for categorization, see Supplementary Methods

Note: presented visually in Fig. 5

### **Supplementary Table S7B: Estimated variation in MIC according to different sources from the random-effects model**

| **Variance Component** | **Test Method** | **Standard Deviation in MIC (95% CI)** |
| --- | --- | --- |
| **Between experiment run (batch)** | CLSI | 0.12 (0.07,0.19) |
|  | EUCAST | 0.20 (0.12,0.32) |
| **Within isolate** | CLSI | 0.38 (0.36,0.40) |
|  | EUCAST | 0.67 (0.64,0.72) |
| **Between isolate** | CLSI | 0.70 (0.64,0.77) |
|  | EUCAST | 0.98 (0.89,1.09) |

Note: Between experiment run: variation due to differences in procedures between experimental runs (batches).

Within isolate: variation intrinsic to isolates between experimental run.

Between isolate: unexplained between isolate variation
